# Label2label: Training a neural network to selectively restore cellular structures in fluorescence microscopy

**DOI:** 10.1101/2020.12.21.423789

**Authors:** Lisa Sophie Kölln, Omar Salem, Jessica Valli, Carsten Gram Hansen, Gail McConnell

## Abstract

Fluorescence microscopy is an essential tool in cell biology to visualise the spatial distribution of proteins that dictates their role in cellular homeostasis, dynamic cellular processes, and dysfunction during disease. However, unspecific binding of the antibodies that are used to label a cellular target often leads to high background signals in the images, decreasing the contrast of a cellular structure of interest. Recently, convolutional neural networks (CNNs) have been successfully employed for denoising and upsampling in fluorescence microscopy, but current image restoration methods cannot correct for background signals originating from the label. Here, we report a new method to train a CNN as content filter for non-specific signals in fluorescence images that does not require a clean benchmark, using dual-labelling to generate the training data. We name this method *label2label* (L2L). In L2L, a CNN is trained with image pairs of two non-identical labels that target the same cellular structure of interest. We show that after L2L training a network restores images not only with reduced image noise but also label-induced unspecific fluorescence signal in images of a variety of cellular structures, resulting in images with enhanced structural contrast. By implementing a multi-scale structural similarity loss function, the performance of the CNN as a content filter is further enhanced, for example, in STED images of caveolae. We show evidence that, for this loss function, sample differences in the training data significantly decrease so-called hallucination effects in the restorations that we otherwise observe when training the CNN with images of the same label. We also assess the performance of a cycle generative adversarial network as a content filter after L2L training with unpaired image data. Lastly, we show that a CNN can be trained to separate structures in superposed fluorescence images of two different cellular targets, allowing multiplex imaging with microscopy setups where the number of excitation sources or detectors is limited.

## Introduction

In recent years, deep learning has increasingly been used for image processing in cell biology [1]. Specifically, convolutional neural networks (CNNs) with, or closely related to, a U-Net architecture [2] are employed for various tasks, from protein detection in transmission microscopy images [3], [4], to image segmentation of single cells [5] or cellular structures such as the nucleus [6]. CNNs have also been used in medical applications, for example, for cell density estimation in corneal endothelial specular microscopy images [7] or tumour classification of urothelial cell carcinoma in pathology [8].

Fluorescence microscopy is a commonly used technique in cell biology to determine the spatial distribution and abundance of target proteins in cells. It relies on the use of highly specific labels to visualise the cellular components of interest. Fluorescent labels can be chemical stains, antibody (AB) labelling or molecular labelling, where cells are genetically altered to express fluorescent proteins [9]. Here, we mainly focus on immunofluorescence (IF) labelling in fixed cells.

Recently, CNNs were employed for content-aware image restoration (CARE) of corrupted fluorescence images. By training a U-Net with image pairs that were generated at different laser powers, or different exposure times, respectively, it was shown that a trained network was able to successfully restore denoised images of cell structures like the cell membrane and the nucleus [2], [10]. The CNN could further be utilised to up-sample images that were detected below the Nyquist sampling frequency [10]. Moreover, generative adversarial networks (GAN) were trained to enhance image resolution in IF microscopy images, using total internal reflection microscopy (TIRF) or stimulated emission depletion (STED) microscopy to acquire the training benchmarks [11], [12].

Both mentioned methods rely on clean benchmark images for the network training which are challenging to acquire in practice. Hence, semi-/unsupervised deep learning-based restoration methods have emerged in IF microscopy. For instance, networks based on the architecture of a cycle generative adversarial network (cycleGAN) [13] were employed to deconvolve fluorescence images of microtubules, using unpaired higher resolution images as reference [14] or simulated low- and high-resolution images [15] for the training. CycleGANs allow the training with unaligned image pairs, addressing the “hallucination problem”, - the introduction of artificial features in the generated images, - when training a classical GAN with unpaired data by implementing an additional training instance in which a second GAN is trained to translate the generated, fake image back to the input [13], [15]. Other examples include *noise2void*, an unsupervised method to remove camera shot noise [16], *DivNoising*, which is based on a variational autoencoder that is trained to restore a distribution of denoised images based on a noise model [17], and *noise2noise* (N2N) [18]. For N2N, a neural network is trained with corrupted image pairs of the same sample. Here, due to the statistical nature of how loss is minimised by the network during training with a deviation-minimising loss function, it was shown that uncorrelated image signals like Gaussian, Poisson or Bernoulli noise are rejected while correlating fluorescence signal is recovered by a network (see ‘Loss Functions for Training a CNN’) [18].

While these image restoration methods enhance the contrast in IF images by upsampling or denoising, all mentioned methods do not correct for inherent background signals in the specimen itself which, in IF microscopy, originate from the cell label or labelling protocol. Unspecific labelling by a stain or antibody binding, as well as internalisation or residue dyes after the specimen preparation, can significantly limit the image contrast of a cell component of interest [19]. The performance of antibodies or stains can vary for different reasons. For instance, epitopes can be altered in the target protein by the fixation step, effectively changing the location or accessibility to the antibody [20]. Also, unspecific antibody binding can be caused by attractive intermolecular interaction such as van der Waals or hydrogen bonding interactions, or by binding to proteins with similar epitopes, which overall results in an underlying cytosolic background signal in images of cells [21].

### Label2label

We propose a new application of deep learning in fluorescence microscopy where a neural network is trained as a content filter of label-induced unspecific cytosolic signals in fluorescence images of distinct cellular structures. We call this method *label2label* (L2L). For L2L, a CNN is trained with image pairs of cells that were dual-labelled for the same distinct cellular structure of interest. L2L utilises the varying performance of antibodies and stains in fluorescence microscopy. We hypothesized that a CNN trained with two images of a cellular structure that originate from two non-identical labels and therefore exhibit sample differences would act as a content filter - where fluorescence signals that systematically vary in the images are rejected, while correlating, cellular structures are restored. Therefore, L2L is different to N2N. In both methods a network is trained without clean benchmark images, but in L2L differences between the training input and benchmark images are not only originating from dynamic image corruptions like noise, but also inherent sample (=label) differences. Consequently, undesired signals originating from cytosolic protein and unspecific binding are retrieved in restorations after N2N training, whereas with L2L we show that a network can be trained as filter for such image content after strategic selection of the training data.

L2L is also different to restoration methods with a *noise2clean* approach such as CARE, where a CNN is trained with images of the same label that were taken under different imaging conditions [10]. While image pairs for *noise2clean* training can be generated with one microscope by, for example, varying exposure time, frame averaging or the sampling density, this approach is rarely necessary in IF microscopy where cell specimens are non-dynamic and exhibit comparably high photon counts. Here, generating the necessary image pairs for the network training is time-consuming and significantly complicated by stage drift resulting in a low benefit-cost ratio. More importantly, as in N2N, background signals originating from the label are still present in the training benchmark, and as such they cannot be corrected by this method.

For L2L training, the images for the training are acquired mostly simultaneously of dual-labelled cells, under near-identical imaging conditions (see ‘Methods: Imaging’). The images obtained using one fluorescent cell label are selected as training input, while the images from the label that yield a higher contrast of the cellular structure are used as training benchmark. We selected the CSBDeep framework for the training that was previously used for CARE of noisy or under-sampled fluorescence images [10]. Moreover, for one dataset, we trained a cycleGAN with unpaired images of the two labels to assess if, in principle, a network can also be trained as content filter using IF images that stem from two separately prepared and imaged cell specimens for the training [13].

To establish and evaluate how our method performs we generated image data for L2L training across four different distinct cellular structures: the actin cytoskeleton, the microtubule network, caveolae and focal adhesions. Actin is a conserved protein in eukaryotic cells that plays a major role in cellular functions like cell migration, cell motility or sustaining the cell shape [22], [23]. Actin exists in two states: as globular (G-) actin in its monomeric form, or polymerised as filamentous (F-) actin [24]. There is evidence that different actin isomers vary in function and localisation; they can appear, for example, in stress fibres, circular bundles, cell-cell contacts or the cell cortex [25]. Microtubules are fundamental cytoskeletal polymeric structures in all eukaryotic cells that, amongst others, play a part in cell transport, cellular signalling via cilia and cell division [26]. The inhibition as well as the promotion of microtubule assembly has been shown to promote mitotic arrest which makes microtubules a prominent target in cancer therapy research [27]. Caveolae are 60-100 nm plasma membrane invaginations composed of heterooligomeric CAVEOLIN and CAVIN protein complexes and abundant in many mammalian cell types [28], [29]. Caveolae are multifunctional organelles that are implicated in transcytosis, lipid homeostasis and cellular signalling. Both CAVEOLIN1 and CAVIN1 are essential for caveolae formation in non-muscle cells [30]. Focal adhesions (FAs) are cellular membrane-associated multi-protein component biomechanical structures. FAs are integral in the ability for most cells to sense and respond to the extracellular matrix and physical changes in the cellular microenvironment, FAs thereby function as central cellular contact points with the extracellular environment, that relay and transmit dynamic external information as internal cellular signalling events [30], [31].

The ability of a CNN to distinguish cellular structures in fluorescence images that was observable in the qualitative results after L2L training in this work was further assessed by training a CNN to separate a nuclear and a plasma membrane marker in superposed IF images of cells that were labelled with a nuclear marker and an antibody against the plasma membrane protein CD44 (see last results section).

### Loss functions for training a convolutional neural network

We used different loss functions to train a CNN with the aim to restore cell images with enhanced structural contrast. A CNN, which can be described as a function *g*_*θ*_ with all its model parameters *θ*, is trained to minimise the error between two images based on a loss function *L*:

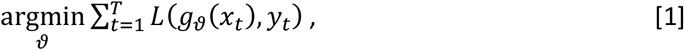

with *g*_*θ*_(*x*_*t*_) as the predicted image for the input *x*_*t*_, *y*_*t*_ as the benchmark and *T* as the number of input-benchmark image pairs that are used for the training [18].

The most commonly used deviation-minimising estimators are the least absolute deviation loss function *L*_*1*_ (or *LAD*) and the least square deviation loss function *L*_*2*_ (or *LSD*) [18], [32]:

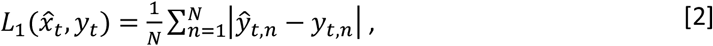

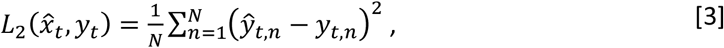

where *ŷ*_*t*_ (=*g*_*ϑ*_(*X*_*t*_)) is the predicted image and *N* is the number of pixels in the image.

Since *L*_*2*_ is minimal if it equals the mean value of the observations, it was previously used for N2N training in cases where the image corruption is, for example, Gaussian noise that, on average, is zero [18]. *L*_*1*_ is the loss function of the CSBDeep framework in default configuration (for non-probabilistic training). With both, *L*_*1*_ and *L*_*2*_, the network calculates the average loss on a pixel-to-pixel basis which limits the ability of the network to consider 2-dimensional correlating signal of structures in an image. As a result, predicted images from a CNN trained with *L*_*1*_ or *L*_*2*_ are often of low quality for a human observer [32], [33].

A multi-scale structural similarity (*MS-SSIM*) index was proposed to compare structural signal between two images that takes the properties of the human visual system into account [34], [35]. The *M*-scale *SSIM* index for a pixel *p* is calculated as follows, after making assumptions as outlined in [35]:

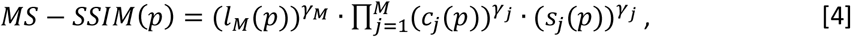

with the exponent *γ*_*j*_ as the weight for the individual scale *j* 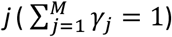, and measures that compare the luminance (*l*_*M*_), the contrast (*c*_*j*_) and structure (*s*_*j*_) of two images. These measures are functions of the local statistics: the sample mean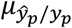, the sample variance 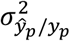 and the sample correlation coefficient 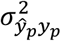 between two image patches *ŷ*_*p*_ and *y*_*p*_ that are the local neighbourhood of a centre pixel *p* [34]. They are described as:

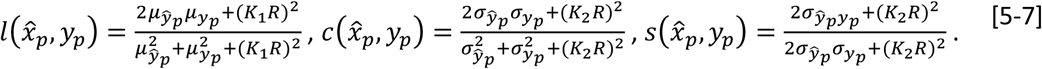

Here, *K*_*1*_ and *K*_2_ are small constants, *R* is the dynamic range of the image and a Gaussian weighting function is used to calculate the local statistics of a pixel in the two images (as in [34]). To calculate the *MS-SSIM* index for *M* > 1, a low-pass filter is applied to the image patches after each iteration, followed by down-sampling by a factor of 2. This approach makes the MS-SSIM index sensitive to differing viewing conditions, such as differing perceived resolution from the point of observation. In [35], the weights *γ*_*j*_ were experimentally determined as (0.0448, 0.2856, 0.3001, 0.2363, 0.1333) to calculate the *5S-SSIM* index (*M*=5) for a representative image example.

The *MS-SSIM* index exhibits values between (−1, 1) where 0 implies no structural similarity and −1/1 a negative/positive correlation between two images. Since a network aims at minimising loss during training, for a multi-scale SSIM loss function (*L*_*MS-SSIM*_) follows [32]:

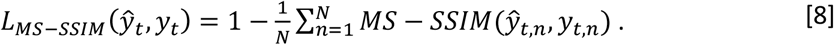

We hypothesized that a CNN would restore images with higher structural contrast after L2L training with a MS-SSIM loss function compared to when using a “classical” *L*_*1*_. To test this hypothesis, we trained a CNN using a *L*_*1*_, *L*_*SSIM*_ (*M*=1), *L*_*3S-SSIM*_ (*M*=3) and a *L*_*5S-SSIM*_ (*M*=5), respectively. The results of the trainings were compared to the denoising method N2N. For N2N training, the same pre-processing steps and network settings were applied, but two noise realisations of the same label were used as training input and benchmark instead that were acquired through sequential imaging. We wondered if using a *L*_*MS-SSIM*_ instead of a *L*_*1*_ for N2N training, would train a CNN to restore images not only with reduced image noise but also increased contrast of the cellular structure. The aim of this work is to compare two deep learning-based restoration methods that improve the contrast of cellular structures in fluorescence images, do not require clean benchmark data and whose requirements for the training data generation are feasible in standard IF microscopy. We show that by introducing systematic sample differences in the training data a CNN can be successfully trained to reject not only image noise but also diffuse cytosolic signals in IF images that are label-dependent and, in practice, decrease the image contrast of a target structure.

## Results

### Label2label to filter out de-polymerised β-actin in images of HeLa cells

To study the performance of a CNN after L2L training with images of the actin cytoskeleton, image pairs of HeLa cells (*N*=68) were generated that were fixed 6 hours after plating, and dual-labelled with the monoclonal anti-β-actin antibody AC-15 and a phalloidin stain (see ‘Methods: Sample preparation for IF microscopy’ and ‘Methods: Imaging’). **Figure 1A** shows an example confocal image pair of the fixed cell specimen. While the phalloidin stain labels almost exclusively the actin filaments, images of the monoclonal anti-β-actin exhibit a high underlying fluorescence signal in the cell body, likely caused by unspecific binding and/or binding to cytosolic protein that result in high intensity punctate regions in the cytoplasm. Notably, a “cleaner”, 20-frame average image of AC-15 exhibits less image noise, but non-filamentous background signal is still present throughout the cell body (see **Figure 1C**). The difference in image contrast between both labels is quantifiable by calculating the mean Michelson and RMS contrast for the images of each label (see [36]). To calculate these, we applied a Gaussian filter (*sigma*=2) to the images, normalised them to their 1^st^ and 99.9^th^ percentile, and derived the sample intensities using the assumption that the 10% brightest pixels in the image represented the sample. For the cell images of the antibody AC-15 and the phalloidin stain, the mean Michelson contrast values of 0.44±0.08 and 0.96±0.04, as well as mean RMS contrast values of 0.15±0.03 and 0.19±0.01 were calculated. Consequently, images of AC-15 were used as training input and images of the phalloidin stain as benchmark for L2L training. For N2N, two noise realisations of AC-15 were used as training data which were acquired through sequential imaging.

**Figure 1.**
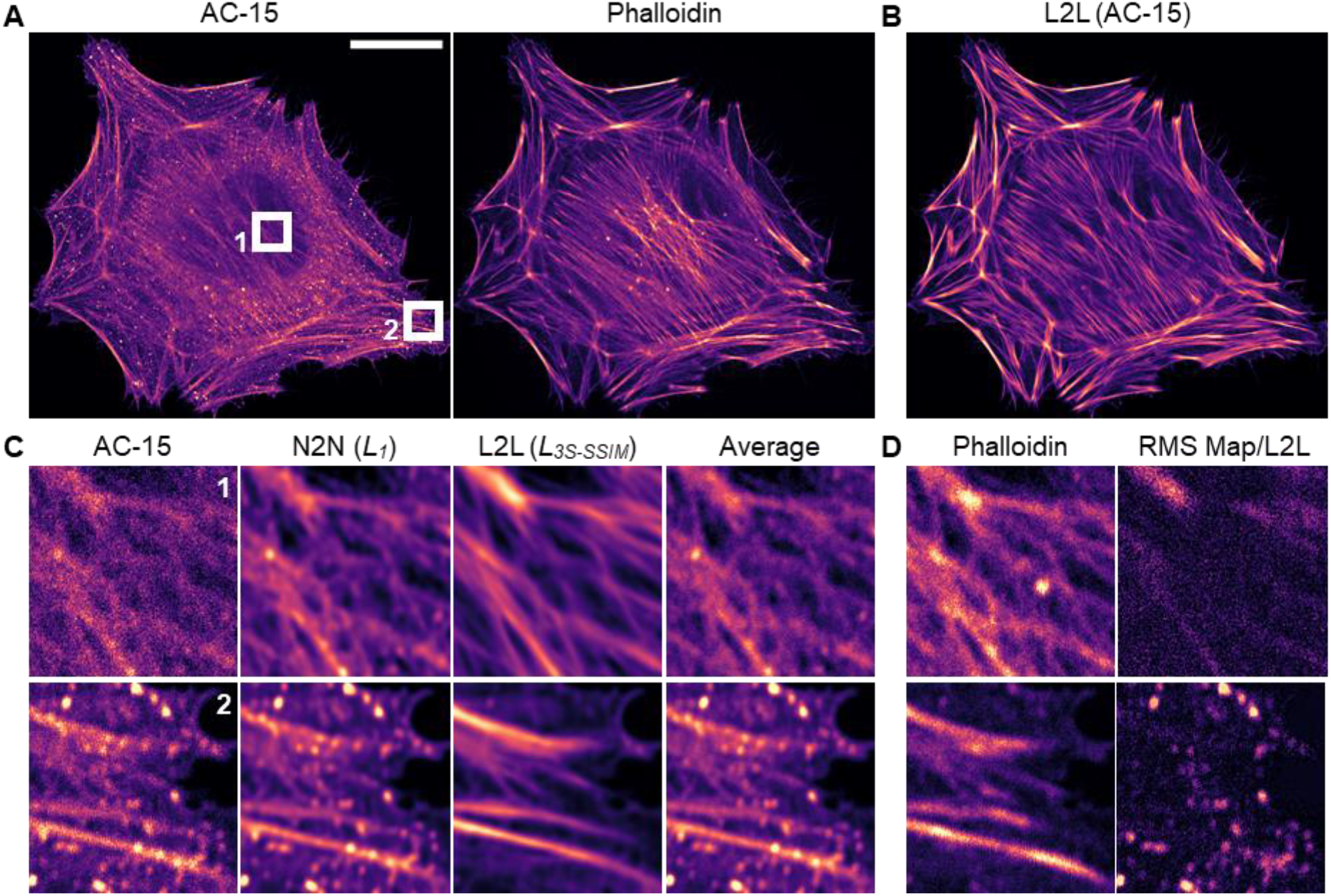
Qualitative label2label (L2L) and noise2noise (N2N) results for images of actin. **(A)** Confocal image pair of a HeLa cell that was fixed, permeabilized and dual-labelled with the anti-β-actin antibody AC-15 and a phalloidin stain which was excluded from the CNN training (scale bar = 20 µm). (**B**) Reconstructed image of AC-15 by a CNN after L2L training with images of AC-15 and phalloidin as training input and benchmark, respectively, using a L_3S-SSIM_ loss function. (**C**) Original and processed images of AC-15 for two ROIs (6 µm x 6 µm). From left to right: raw image, restored images after N2N and L2L training with a L_1_ and L_3S-SSIM_ loss function, respectively, and a 20-frame average. (**D**) The corresponding phalloidin image and the RMS map between the raw and the predicted image of AC-15 of the network after L2L training.

**Figure 1B** shows the restoration of an image of AC-15 in a HeLa cell (see **Figure 1A** (*left*)) by a CNN after L2L training with a 3-scale structural similarity loss function (*L*_*3S-SSIM*_) (see ‘Methods: Training the CNN’). The CNN reduces cytoplasmic signal throughout the cell body, while the relative signal of filamentous actin labelled with AC-15 is increased in the restored image. In **Figure 1C**, for two regions of interest (ROIs), the original cell image of AC-15 as well as the predicted images of a CNN after N2N and L2L training with a *L*_*1*_ and *L*_*3S-SSIM*_, respectively, are shown. While noise is reduced in the restored images after N2N training, resulting in an optically similar result to a 20-frame average, the relative contrast between the actin filaments and the background signal that originates from depolymerised protein is not enhanced. In the L2L results, however, not only image noise is removed, but also the contrast of filamentous signal is clearly enhanced, even compared to the training benchmark (phalloidin) (see **Figure 1D** (*left*)). Here, high intensity punctate regions originating from unspecific binding or depolymerised actin are selectively filtered out by the CNN as evident in the RMS maps between the raw images of AC-15 and the L2L results (see **Figure 1D** (*right*)).

In **Supplementary Information (SI) Figure 1**, the qualitative results for N2N and L2L are shown for AC-15 after training with different loss functions (*L*_*1*_, *L*_*5S-SSIM*_, *L*_*3S-SSIM*_, *L*_*SSIM*_), including the corresponding images of the training benchmark (phalloidin) for L2L training (see panel C) and 20-frame average images of both labels (see panel A+C) to better assess the results. For both methods, using a least absolute deviation loss function (*L*_*1*_) for the training leads in comparison to more conservative predictions, where, in L2L, non-filamentous signal is filtered out by the network, but actin filaments appear relatively blurry. On the other hand, structures in restored images after training the CNN with a single-scale SSIM loss function (*L*_*SSIM*_) appear sharp with a higher likelihood of appearing erroneous predictions. These hallucination effects are to a greater extent noticeable in the results for N2N compared to L2L. Here, punctuate regions in the images of AC-15 appear sharp in the restorations. Also, using a *L*_*SSIM*_, filaments that appear with low contrast in the raw images of AC-15 are restored with intensity fluctuations along the structure after N2N training, which is not observed for L2L.

To further evaluate the performance of the network after L2L training, the average peak signal-to-noise-ratio (*PSNR*), the normalised root-mean square error (*NRMSE*), and the average *5S-SSIM, 3S-SSIM* and *SSIM* indices were calculated for the raw or predicted images of AC-15 and the corresponding images of phalloidin, dependent on the training loss function. These were calculated using the validation image patches that were excluded from the training data (see ‘Methods: Training the CNN’). All calculated metrics indicate that the correlation between the predicted images for AC-15 after L2L training and the benchmark (phalloidin) is increased compared to the original image (see **Table 1A**). Notably, the network performance increases after replacing a standard *L*_*1*_ with a MS-SSIM loss function, with the *L*_*5S-SSIM*_ narrowly yielding the best *PSNR* and *NRMSE*.

**Table 1.**
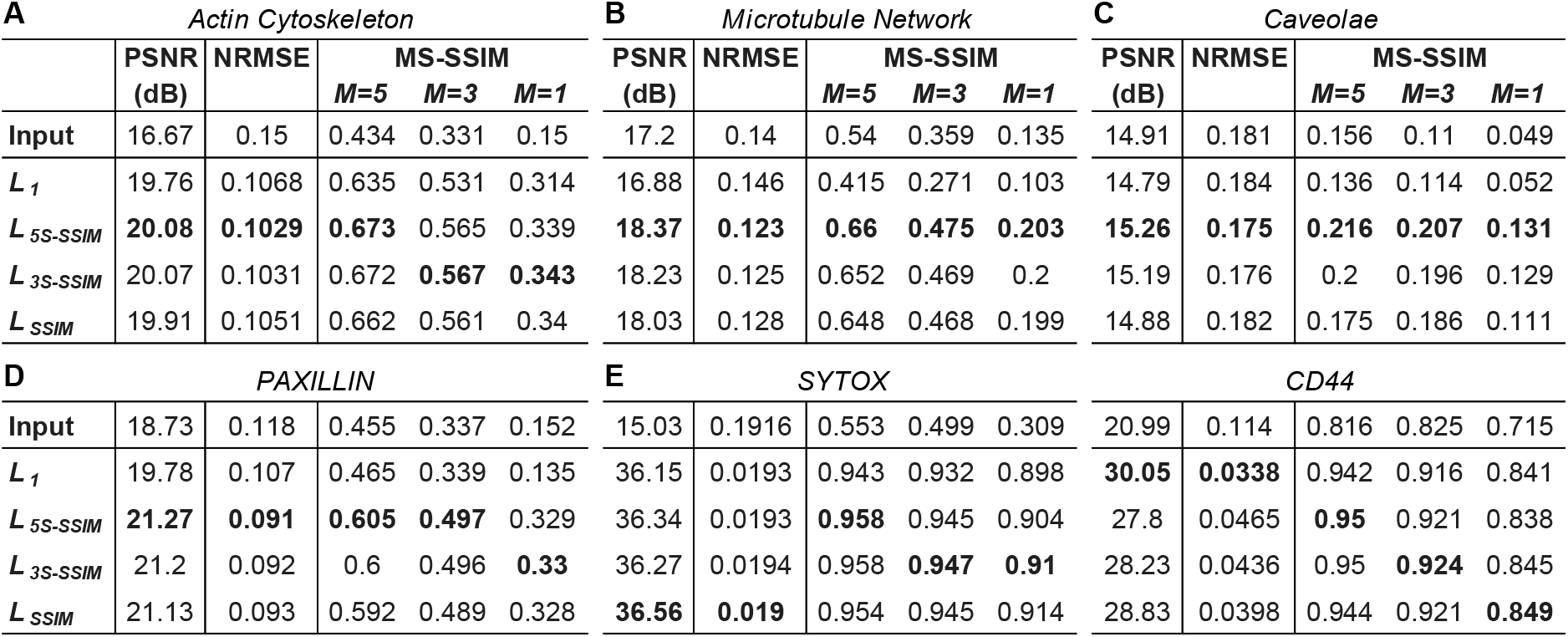
Loss function-dependent evaluation of label2label (L2L) for images of different cellular structures. Evaluation after training a CNN with **(A)** image pairs of HeLas that were dual-labelled with an anti-actin antibody (input) and a phalloidin stain (benchmark), (**B**) of MeT5As that were labelled with the anti-α-tubulin antibodies DM1A (input) and YOL1/34 (benchmark), **(C)** of MeT5As that were labelled with antibodies against CAVIN-1 (D1P6W) (input) and CAVEOLIN-1 (4H312) (benchmark), and (**D**) of MeT5A, HeLa and U2OS cells that were dual-labelled with the anti-PXN antibodies 5H11 (input) and Y113 (benchmark). In (**E**), the results are shown after training a CNN to separate markers in superposed images of cells that were labelled with the nuclear marker SYTOX and an anti-CD44 antibody. Average metrics were calculated between the respective training inputs or restored images of the input, respectively, after training the network with a L_1_, L_5S-SSIM_, L_3S-SSIM_ or L_SSIM_ loss function, and the corresponding training benchmarks, using the image patch pairs that were excluded from the training for the validation (see **Table 2**). Metrics for the caveolae dataset were calculated after applying a Gaussian filter (sigma=1.5) due to high image noise in the benchmark images. The best value for each column is shown in bold.

### Label2label to enhance the structural contrast in images of the microtubule network and caveolae

To further study how L2L training impacts the contrast of distinct cellular structures in the restorations of a trained network, fluorescence image pairs were acquired of the microtubule network that has a distinct branched, spatial distribution in cells, and caveolae that are 60-100 nm large invaginations in the plasma membrane [29], [37]. For the former, fixed MeT5A cells (*N*=51) were dual-labelled with two monoclonal antibodies against α-tubulin (DM1A raised in mouse and YOL1/34 raised in rat) and confocal image pairs were generated. To train a CNN to increase image contrast of caveolae structures, STED image pairs of caveolae (*N*=60) were acquired of fixed MeT5A cells that were dual-labelled for the two essential caveolae components CAVIN-1 (D1P6W) and CAVEOLIN-1 (4H312) [28]. For the tubulin dataset, we calculated mean Michelson contrast values of 0.45±0.14/0.9±0.05 and mean RMS contrast values of 0.13±0.03/0.17±0.02 for DM1A/YOL1/34; for the caveolae dataset, these values were 0.27±0.05/0.98±0.02 and 0.13±0.02/0.22±0.02 for D1P6W/4H312 (see previous section; for the caveolae dataset, the 1% brightest pixels in the image were regarded as sample). Hence, for L2L training, images of DM1A and D1P6W were used as input, images of YOL1/34 and 4H312 as benchmark to train a CNN as content filter for images of microtubules and caveolae, respectively.

In the cell images of YOL1/34 (tubulin) we observed intensity fluctuations along the microtubules that might originate from selective binding to specific epitopes of polymerised tubulin or in-homogeneous binding, which was not observable for the clone DM1A (see high-frame average images in **SI Figure 2C**). The microtubules appeared overall sharper in the cell images of YOL1/34 compared to DM1A, with a lower, observable “haze” in the cytoplasm. This haze was likely caused by unspecific binding, binding to cytosolic tubulin and/or out-of-focus signal – where the optical resolution in images of DM1A labelled with the secondary antibody Alexa Fluor 633 was lower than in images of YOL1/34 which was conjugated to Alexa Fluor 488.

In **Figure 2A**, the results for the tubulin dataset after N2N and L2L training with a *L*_*1*_ and *L*_*3S-SSIM*_ for two ROIs are displayed. The top row shows the results for a representative training image pair; in the bottom, an image pair is shown that was acquired for the same sample, but with a different imaging setup that allowed confocal and STED imaging (see ‘Methods: Imaging’). STED images with an enhanced resolution of circa a factor 3 were acquired to allow a better assessment of the restoration performance of the CNN for this sample, since microtubules are insufficiently resolved in confocal microscopy, especially if densely packed [37]. Structural contrast in the images of tubulin is increasingly enhanced between the restorations of a network after “classical” N2N training with a *L*_*1*_ and L2L training with a *L*_*3S-SSIM*_ (see **Figure 2A**). Both methods restore images with higher contrast of the microtubule structure compared to a 20-frame average image (see **Figure 2A** (*top*) and **SI Figure 2**). The restorations for the cell image of DM1A that was acquired with different imaging settings than the training input exhibits a slightly less homogeneous intensity distribution along the microtubules, likely due to the differing noise level (compare **Figure 2A** (*left*)). For both, N2N and L2L, the restoration success is dependent on the tubule density in the image. The closer microtubules are packed in the cell, the less likely is the successful recovery of separate structure by the CNN as evident by comparing the restoration of the confocal image of DM1A with its STED image (see **Figure 2A** (*bottom*)). Notably, the intensity fluctuations along the cellular structure in the training benchmark YOL1/34 (see **Figure 2B**) do not result in artefacts in the restored images of DM1A after L2L training. Further restorations of representative training inputs are shown in **SI Figure 2** after N2N and L2L training, respectively, using a *L*_*1*_, *L*_*5S-SSIM*_, *L*_*3S-SSIM*_ or *L*_*SSIM*_. A loss function-dependent trend for both methods is observable: using a MS-SSIM loss function with decreasing scale instead of a *L*_*1*_ for the training enhances the contrast and sharpness of microtubules in the restorations, with L2L training leading to a higher contrast of the cellular structure than N2N (see **SI Figure 2**).

**Figure 2.**
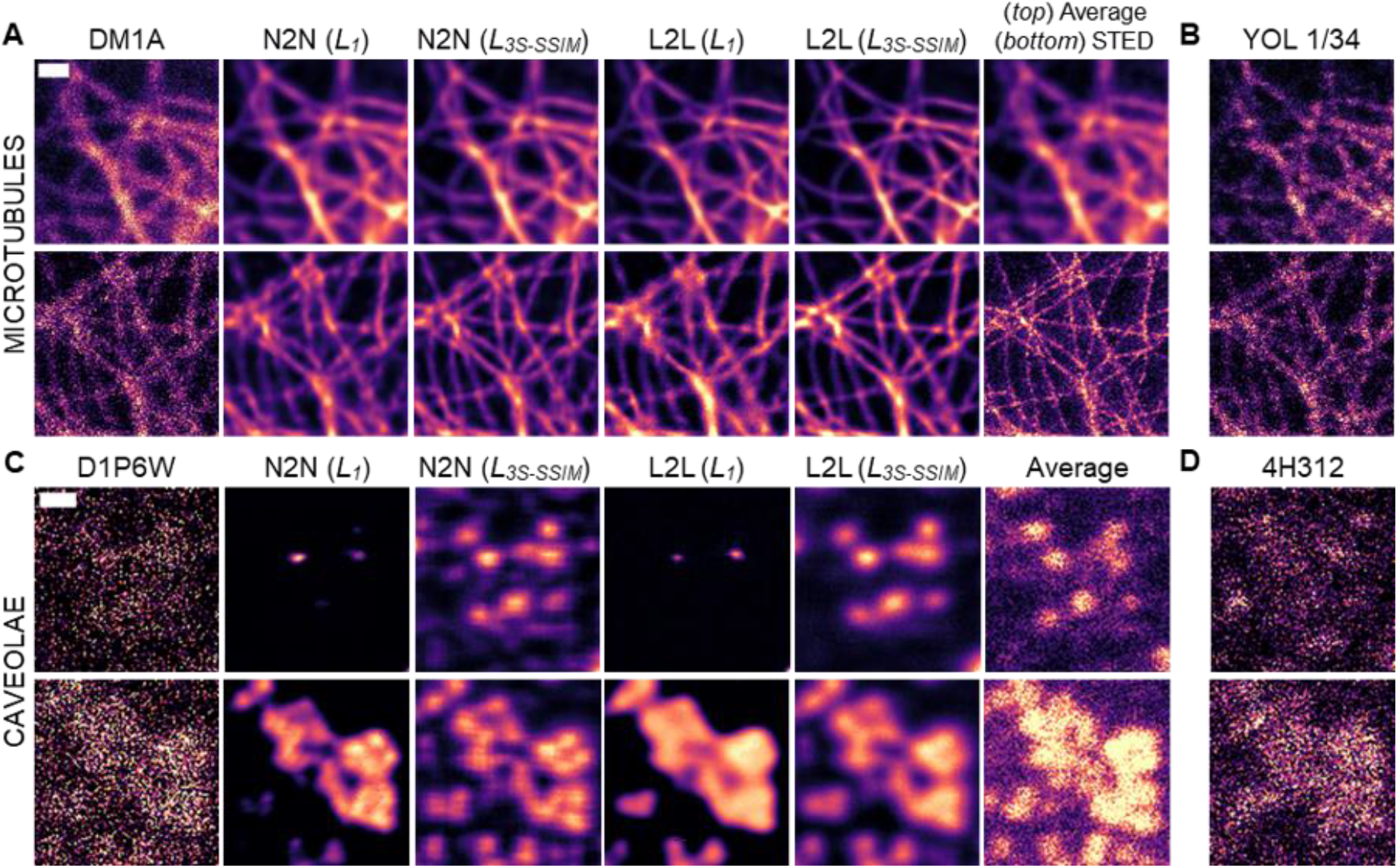
Loss function-dependent label2label (L2L) and noise2noise (N2N) results for images of the microtubule network and caveolae. Confocal images of MeT5A cells that were dual-labelled with the (**A**) anti-tubulin antibodies DM1A and (**B**) YOL1/34 (scale bar = 1 µm), and STED images of MeT5A cells that were dual-labelled with (**C**) the anti-CAVIN-1 antibody D1P6W and (**D**) the anti-CAVEOLIN-1 antibody 4H312 (scale bar = 200 nm). **(A, C)** From left to right: raw image of a representative training input for the trainings, reconstructed images after N2N and L2L training with a L_1_ or L_3S-SSIM_ loss function and a corresponding 20-frame average or high resolution STED image. In **(B, D)**, representative training benchmarks for L2L training are displayed. Shown images were excluded from the network training.

The STED image pairs which were acquired of fixed cells that were dual-labelled for the two essential caveolae proteins CAVIN-1 (D1P6W) and CAVEOLIN-1 (4H312) exhibit a low signal-to-noise ratio, further amplified by the pre-processing step that was undertaken to prevent artefacts we observed when training with the nearly binary, raw data (see ‘Methods: Data augmentation and pre-processing’; see **Figure 2C-D**). For some image pairs, we found that the correlation of the fluorescence signal originating from caveolae structures was relatively low between the training input (D1P6W) and benchmark (4H312) (see **SI Figure 3A+C**). This is also indicated by the calculated mean *PSNR, NRMSE* and *MS-SSIM* indices for both images that showed the lowest correlation of the training data for all datasets that were acquired for L2L training in this work (compare **Table 1A-D**). It posed the question if a CNN would be prone to introduce artificial structure after L2L training with this dataset.

We found that the CNN performance after N2N and L2L training is highly dependent on the utilised loss function (see **Figure 2C**). Using a *L*_*1*_, for both training modalities, a trained CNN only restores high signal-to-noise regions in the images (compare **Figure 2C** (*top*) and (*bottom*)). Using a *L*_*3S-SSIM*_, however, the recovery of structural signal by the CNN is much enhanced for N2N and L2L. Here, background signal in the image, likely originating from unspecific labelling, is filtered-out after L2L training, while, with N2N, the CNN recovers weak structure-like signal in such image areas (**Figure 2C** (*top*)). Also, weak stripe-like artefacts appear in the N2N results (see also **SI Figure 3B**). These might originate from small non-dynamic image corruptions from the imaging setup that are present in sequential images of D1P6W, and that are picked up by a loss function that is sensitive to structural similarity of images (*L*_*MS-SSIM*_). The sample differences in the training data for L2L cause the CNN to restore caveolae structures with slightly lower sharpness compared to N2N but it does not lead to observable hallucinations effects. Notably, the level of sharpness as observed in the N2N result is not verifiable in the 20-frame average image (see **Figure 2C** (*bottom*)). The introduction of artefacts by a CNN after N2N training with a MS-SSIM loss function is also clearly visible in **SI Figure 3**, where restored images are shown after training a CNN for N2N and L2L with a *L*_*1*_, *L*_*5S-SSIM*_, *L*_*3S-SSIM*_ or *L*_*SSIM*_. While low signal-to-noise regions in the image are barely restored by the CNN using a *L*_*1*_, using a MS-SSIM loss function for N2N training, the network learns to restore background signal from the sample as caveolae-like structure. This is not observed for L2L. Here, using, for example, a *L*_*3S-SSIM*_ for the training, the network restores the noisy image by selectively recovering caveolae structure and filtering out background signal that stems from the sample, resulting in much enhanced results - also compared to the 20-frame average images of both labels (see **Figure 2** and **SI Figure 3**).

Again, the network performance was evaluated after L2L training with different loss functions by calculating the PSNR, NRMSE and different MS-SSIM indices (see **Table 1B-C**). For both datasets, the calculated metrics indicate that the correlation between reconstructed and benchmark image is highest after training with a *L*_*5S-SSIM*_. For the tubulin dataset, a decrease in the correlation between the restored image of DM1A and the benchmark (YOL1/34) after training with a *L*_*1*_ is observed.

### Training networks to reduce cytosolic content in images of PAXILLIN with paired and unpaired images

We also trained a CNN with image pairs of two non-identical labels against the focal adhesion (FA) protein PAXILLIN (PXN) [31], with the aim to reduce fluorescence signal in the images that stems from cytosolic protein. Image pairs of fixed MeT5A (*N*=47), HeLa (*N*=17) and U2OS (*N*=13) cells were acquired that were dual-labelled with the monoclonal anti-PXN antibodies 5H11 and Y113. As expected, the raw IF images of both antibodies show correlating FA structures in the cells, but also a diffuse signal throughout the cell body that differs in relative intensity to the FA signal between the two labels (compare **Figure 3A** (*left*) and **Figure 3B**). For all cell lines we observed the same trend; the images of the antibody Y113 exhibit a higher FA-to-cytosolic signal ratio compared to 5H11. To determine if the cytoplasmic signal originates from clone-dependent binding to protein in the cytosol or unspecific binding, shRNA mediated *PXN* knockdown MeT5A cells were generated (see ‘Methods: Generation and verification of knock down PAXILLIN MeT5A cells’). The knockdown was confirmed via a qPCR analysis that shows a knockdown efficiency of >70% on mRNA level, and substantially lower protein levels were confirmed by Western blotting (see **SI Figure 4A-B**). For both antibodies, the IF signal of PXN was overall reduced in images of fixed shPXN cells in the FAs and in the cytoplasm (see **SI Figure 4C**). Hence, the cytoplasmic signal that is observed in cell images of both clones is largely not caused by unspecific binding. Instead, the different relative cytosolic content in cell images of 5H11 and Y113 indicates a differing accessibility of the respective binding sites for both clones for protein that is cytosolic or bound to FAs, and that cytosolic PXN likely functions as a readily available replenishable buffer for FA localised PXN.

**Figure 3.**
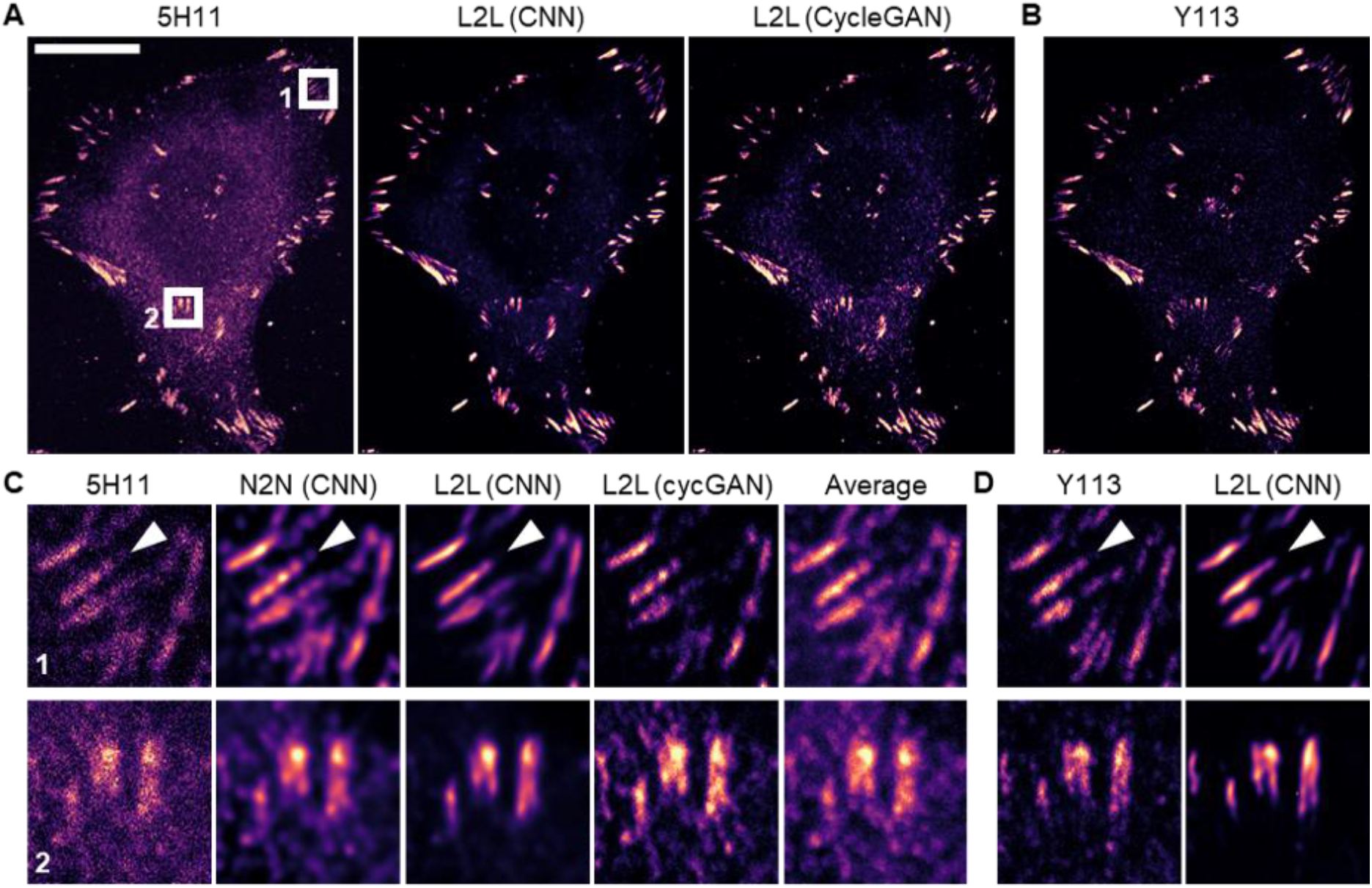
Network architecture-dependent label2label (L2L) results for images of *PAXILLIN*. Confocal images of a HeLa cell that was dual-labelled with the two anti-PAXILLIN antibodies 5H11 and Y113 which were used as training input and benchmark for L2L training. **(A)** From left to right: raw image of 5H11, the restored images of a CNN after L2L training with paired images and of a cycleGAN that was trained with unpaired images, and **(B)** the corresponding image of Y113 (scale bar = 20 µm). **(C)** Predicted images for two ROIs (6 µm x 6 µm). From left to right: input (5H11), the restored images of a CNN after N2N and L2L training with a L_1_, the restored image of a cycleGAN, and a 20-frame average. **(D)** Corresponding benchmark images (Y113) for L2L training and its prediction by a CNN after L2L training as outlined above. A network that was trained with L2L data in-paints focal adhesions (see white arrows) and filters out cytosolic protein (see bottom ROI) for both, the training input and benchmark. Shown images were excluded from the network training.

All cell lines were prepared and imaged under the same conditions, and were used indistinguishable to generate the training data (see ‘Methods: Training the CNN’). For the cell images of 5H11/Y113 (*N*=77), mean Michelson contrast values of 0.57±0.07/0.99±0.01, and mean RMS contrast values of 0.15±0.01/0.18±0.02 were calculated. Consequently, cell images of 5H11 were used as training input and images of Y113 as benchmark to train a network to decrease cytosolic signal in images of focal adhesions. To test if an artificial network can also be trained as content filter with unpaired L2L data, a cycleGAN was trained and its performance was compared to results obtained after training the CNN (see ‘Methods: Training a CycleGAN’).

In **Figure 3A**, the predicted images of a CNN (using a *L*_*1*_) and a CycleGAN for a cell image of 5H11 are shown after L2L training with aligned and unaligned images, respectively. Both networks learn to reduce cytosolic content in images of 5H11 and selectively recover FA structure, but the CNN outperforms the CycleGAN for this task. While the trained cycleGAN acts as content filter of cytosolic signal, high intensity areas in the cell cytoplasm that do not stem from FAs are, in part, still present in the predicted images with a tendency to translate intensity fluctuations in the input as weak structure (**Figure 3C** (*bottom*)). A similar effect is observed after training a CNN for N2N, using a MS-SSIM loss function (see **SI Figure 5B**). Here, the CNN learns to artificially accentuate intensity fluctuations in the cytosol in restored images which become more pronounced with a decreasing number of iterations (*M*), leading to very different qualitative N2N results after using a *L*_*1*_ and a *L*_*SSIM*_. These artefacts are much less pronounced in the L2L results (see **SI Figure 5B**). However, contrary to the results of the evaluation (see **Table 1D**) that, again, indicate the highest network performance after using a *L*_*5S-SSIM*_, subjectively, the *L*_*1*_ loss function was best suited to train a CNN as filter for cytosolic signal for this dataset.

After L2L training with a *L*_*1*_, cytosolic signal is clearly reduced in the predicted images compared to the high average image or N2N result, resulting in images with increased contrast of the FAs (see **Figure 3C**). In addition, the trained CNN in-paints focal adhesion structures that are inhomogeneously labelled (see white arrows in **Figure 3C**). Also, we found that the CNN restored images of cells labelled with Y113 with decreased cytosolic content as well, although these images were used as benchmark, not input for the training (see **Figure 3D**).

### Training a CNN to separate cellular structures in superposed images

Lastly, the ability of a CNN to filter-out features in fluorescence images based on their spatial intensity distribution was tested by training a network to separate superposed confocal images of two different cellular targets. For that, fixed MeT5A cells were stained with the nuclear marker SYTOX Green and labelled with an antibody against CD44, a plasma membrane protein. Then, image pairs (*N*=58) were acquired of both markers (superposed) and of each marker seperately (see ‘Methods: Imaging’). The spatial distribution in the cell of both targets partly overlapped in the images, but the targets were structurally distinguishable - with the nucleus appearing with high intensity in the cell centre and CD44 distributed at the periphery of the cell (see **Figure 4A**). The CNN was trained with the superposed image of both structures as training input in a two channel image, and the separate cell images of SYTOX and CD44, respectively, as training benchmark (see ‘Methods: Training the CNN’). The qualitative results in **Figure 4** for an example image pair show that a CNN seperates the nucleus and plasma membrane marker in the input image successfully after the training. Noticably, similar to N2N, image jitter and noise that are present in the input image are removed by the network in the restorations (see **Figure 4B**). However, the CNN slightly blurs structure in the restorations and overlapping structures in the nuclear area are mostly but not fully recovered in the images of CD44. A loss function-dependent evaluation of the network performance showed no clear trend as to which loss function was best suited to train the CNN as separator of the two structures (see **Table 1E**).

**Figure 4.**
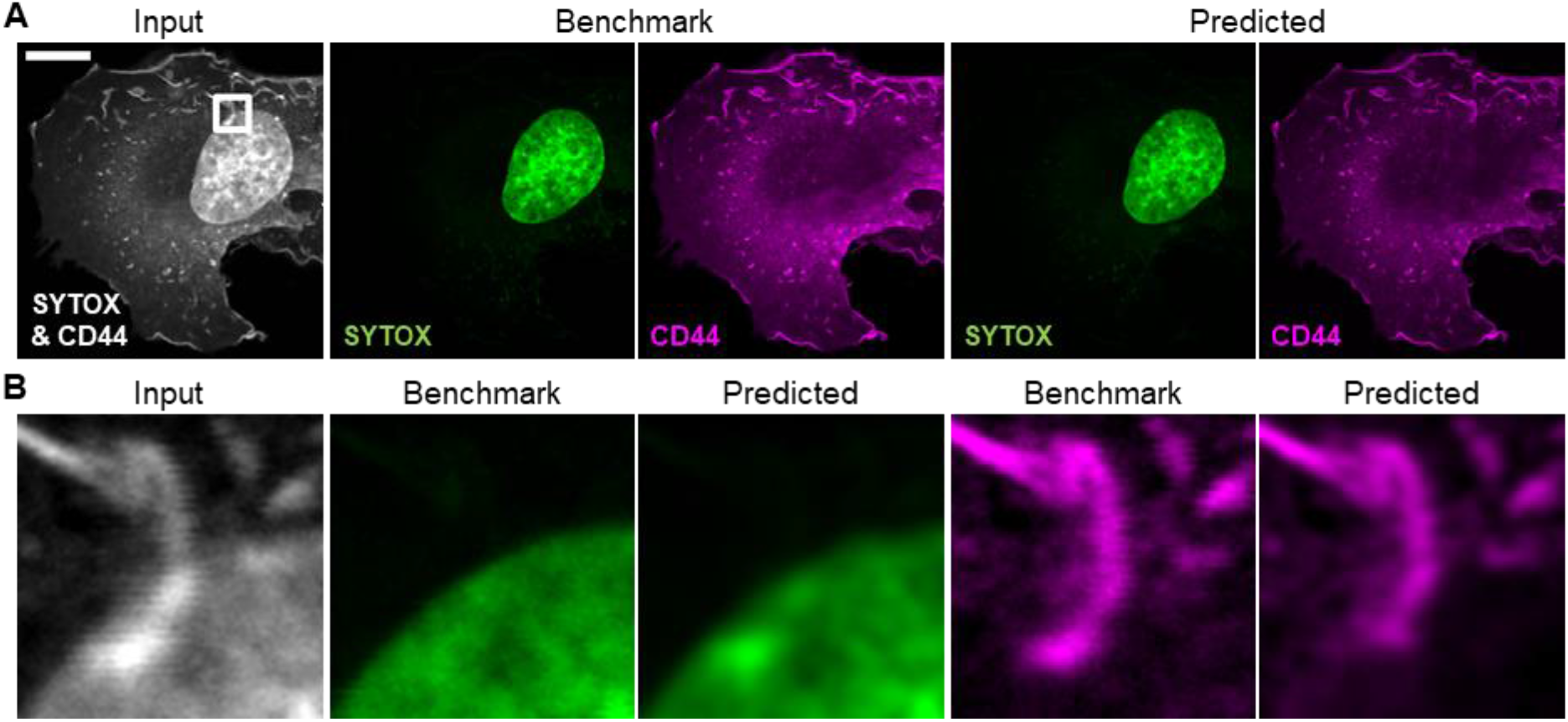
Qualitative results after training a CNN to separate cellular structures in superposed images of a nuclear stain and an antibody against a plasma membrane protein. (**A**) Training input and benchmark images of a MeT5A cell that was dual-labelled with the nuclear stain SYTOX Green and an anti-CD44 antibody, and corresponding reconstructions after training a CNN with a L_3S-SSIM_ (scale bar = 10 µm). The images for training the CNN were obtained via sequential imaging, using different excitation wavelengths. (**B**) Qualitative result for a ROI (5 µm x 5 µm). Prediction success is dependent on the level of superposition of both labels. Structures appear slightly blurry in the restorations compared to the benchmark, but image noise and jitter are reduced. The shown images were excluded from the training.

### Testing the robustness of label2label

The robustness of L2L training was evaluated for the actin (*N*_*tot*_=68), tubulin (*N*_*tot*_=51), caveolae (*N*_*tot*_=60) and PAXILLIN (*N*_*tot*_=77) dataset by studying how much the network performance is impacted by specific image pairs that are used to generate the training data. For that, repeated cross-validations were conducted with randomly selected image pairs *N* from the total dataset (*N*_*tot*_). For the actin and PAXILLIN dataset, we conducted 8-fold cross-validations with *N* = 8, 16, 32 and 64; for the tubulin and caveolae dataset, we conducted 10-fold cross-validations with *N* = 10, 30 and 50. We increased the number of repetitions for small *N*, and the epoch number for the training was linearly adapted to *N* to prevent overfitting (see ‘Methods: Training the CNN’, **Table 3**). The difference between this approach and the default validation in the CSBDeep framework is that in the latter the validation is conducted via a train/test split of the training data which is generated from all raw image pairs. This way, image patches generated from a specific image pair appear in both, the test (validation) and the training data, making it impossible to assess how much the network performance depends on if a specific image pair is used for the training.

The mean relative change of the *NRMSE* and *5S-SSIM* index between the input and the restored image (both versus the reference (=training benchmark)) are shown in **Figure 5**. Each data point represents the mean value of a cross-validation dependent on *N*. A Gaussian filter (*sigma*=1.5) was applied to images of caveolae prior to the calculations since noise levels were so high in the raw input images that no trend was observable for unprocessed data. The results for nearly all datasets show a similar trend: even when using a small *N*, the structural similarity to the benchmark increases for restored images compared to the input. However, the calculated *ΔNRMSE* and *Δ5S-SSIM* indices are dependent on the particular images that are used for the training – indicated by a wide distribution for small *N*. This could indicate that certain acquired image data is better suited to train the network, or easier to predict for the network. As expected, on average, the highest performance was achieved using a high *N*. While the results for *Δ5S-SSIM* for all datasets indicate that with *N* > 30 or 32, respectively, the network performance is relatively consistent, the *ΔNRMSE* improves continuously with increasing *N*. Here, the trend of *ΔNRMSE* deviates for the caveolae dataset for which only slight changes are observed between different *N*, without a clear trend.

**Figure 5.**
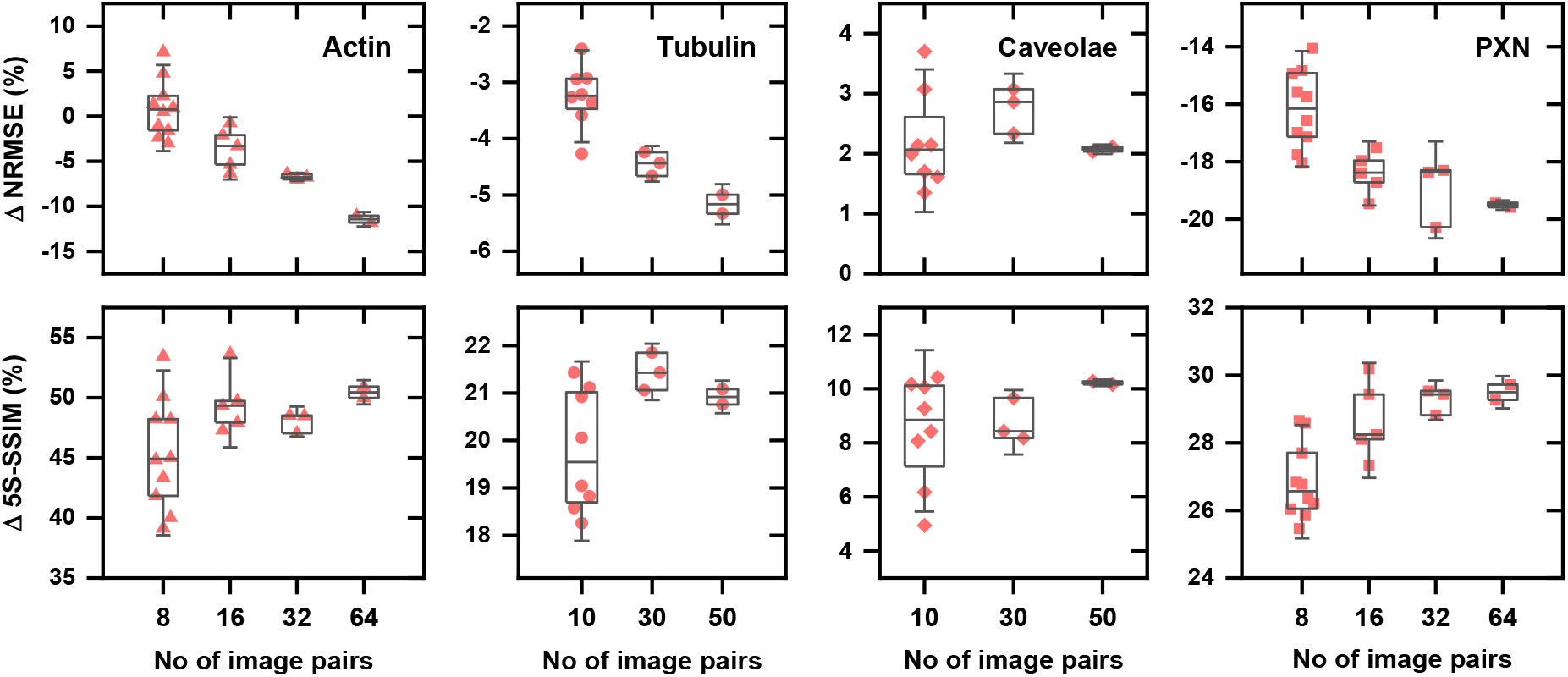
Repeated cross-validation for L2L training. The mean relative change between the initial (input and benchmark) and final (restoration and benchmark) NRMSE and 5-scale SSIM index after L2L training with image pairs of cells that were dual-labelled for the actin cytoskeleton (N=68), tubulin (N=51), caveolae (N=60) or PAXILLIN (N=77), dependent on the number of raw image pairs that were randomly selected from the total dataset for the cross validation. Each data point is the mean value for an 8-fold (actin, PAXILLIN) or 10-fold (tubulin, caveolae) cross-validation that was repeated for small image pair numbers.

## Discussion

We show that a CNN can be successfully trained to filter out unspecific, label-induced fluorescence signals detected in the cell cytoplasm in IF images of cellular structures, using images of two non-identical labels as training data that target the same structure but exhibit systematic image differences. L2L is different to restoration methods that use images of the same label as training data, for example, N2N. While after N2N training a CNN restores all signals originating from the label without distinction, a CNN systematically learns to disregard non-structural signals after L2L training. We found that a network trained with L2L data restored images with high contrast of the target structure, even compared to the training benchmark or its high frame average image (see, for example, **Figure 2A+C**). The trained network could further be utilised to improve images that were not used as input for the training. For instance, while cytosolic content was present in images of cells labelled with Y113 which were used as benchmark to train the network as content filter of cytosolic PXN, it was nearly eradicated in the restoration by a trained CNN (see **Figure 3D**).

The network performance as content filter after L2L training was dependent on the level of correlation between the images of the two labels and the training loss functions. Using a single-scale SSIM loss function for the training increased the likelihood of hallucination effects in restored images after L2L training, especially for the PXN dataset where cytosolic content in the cell body was not entirely without structure and present in both, training input and output. However, a MS-SSIM loss function was better suited to train a network to restore sharp cellular structures. Here, the restorations increasingly (with *M*) converged to results obtained with a *L*_*1*_ (see **SI Figure 1**-**3, 5**). For images of actin, tubulin and caveolae we determined that using a *L*_*3S-SSIM*_ loss function led qualitatively to the best results. For these datasets, the to-filter-out signal deviated sufficiently between the image pairs, allowing the network to clearly distinguish to the cellular structure that correlated in the images of both labels. For the PXN dataset, a *L*_*1*_ was more suited to train a CNN to recover FAs only, likely due to the correlating cytosolic signal in the training images, albeit both at different relative intensity to the FA signal (see **Figure 3A-B**).

Thus far, most deep-learning based image restoration methods have relied on image pairs of the same label with differing resolution or photon counts as training data. For such training data, the N2N results suggest that powerful loss functions like the MS-SSIM loss function are prone to overestimate structural signal, resulting in increased hallucination effects in the restorations (see, for example, N2N results in **SI Figure 3B+5B**). Using a MS-SSIM loss function for N2N training led to a significantly higher occurrence of artefacts compared to L2L, where cytosolic, non-structural content (as present in the training input for the actin, caveolae and PXN dataset) was restored with accentuated, artificial contrast. Only for the tubulin dataset, where background signals in the images of both labels were very unspecific, we observed a clear improvement between the restored images after N2N training with a *L*_*1*_ and a *L*_*3S-SSIM*_, respectively (see **Figure 2A**). The qualitative N2N results for the caveolae dataset indicate that a CNN picks up structure much more efficiently in very noisy images when using a *L*_*3S-SSIM*_ instead of a *L*_*1*_ (see **Figure 2C**). However, using images of the same label, again, caused the network to hallucinate and non-dynamic image corruptions that were likely caused by the imaging system appeared with higher contrast (see **Figure 2C** and **SI Figure 3B**).

Although clean benchmark data are not required for L2L and N2N training, in images of structures that are not sufficiently resolved by the imaging technique, either *a posteriori* knowledge about the structure or clean reference data are required to assess the qualitative performance of a trained CNN. This was especially noticeable in cell images of tubulin, where the fine microtubule network was not fully resolved with confocal microscopy (see **Figure 2A-B**). This led to erroneous predictions in image areas with a dense microtubule network after both, N2N and L2L training.

We found that metrics like the *PSNR* and *NRMSE* were inadequate to forecast which loss function yielded the best restorations by a CNN after L2L training - especially given that the reference image was not clean, showing the cellular structure only. For instance, the calculated metrics after L2L training with different MS-SSIM loss functions barely deviated for all dataset examples while, qualitatively, differences in the restorations were observed (see **Table 1A-D**). While the mean SSIM indices that were calculated for the input and benchmark images indicated scarcely any correlation, with values around 0.05-0.15 for all L2L datasets (see **Table 1A-D**), the initial correlation between the training data was higher according to the calculated *5S-SSIM* indices (by a factor of circa 3). This indicates that a high-scale SSIM index is more suited to detect correlation in fluorescence images of cellular structure than metrics like the *PSNR, NRMSE* or *SSIM* index, likely because the images are the convolved signal of a sample volume rather than the strict 2D depiction of the sample at a specific section. This observation fits to the theory of the MS-SSIM index [35].

As expected, the evaluation of repeated 8/10-fold cross-validations (dependent on the number of raw image pairs that were used to generate the training data) showed that using a high number of image pairs to train the CNN as content filter is advisable (see **Figure 5**). While the mean *PSNR* and *5S-SSIM* index significantly increased even after L2L training with a small number of image pairs, the results were dependent on the selected image pairs. Although the evaluation would be more meaningful if each cross validation had been conducted with unique image pairs instead of image pairs that were randomly selected from a finite dataset, the results indicate that a network converges to the optimal result of a particular dataset during training (see **Figure 5**).

Prior to this work, it was unclear if the difference in the training data would introduce artificial structure in the predicted images of a CNN after L2L training. For instance, in the caveolae dataset, the sample differences between input and output were quite significant (compare **SI Figure 3A+C**), and inhomogeneous labelling of the microtubule structure by the anti-tubulin antibody YOL1/34 (benchmark) was not visible in the training input (DM1A) (compare **SI Figure 2A+C**). Yet, we could not observe that the CNN introduced artificial structure in the restored images after L2L training. Instead, the sample differences proved advantageous when training with a MS-SSIM loss function to significantly decrease the introduction of artificial structure in the restorations that we observed after N2N training. However, an in-painting effect was observed for L2L, primarily in restorations of AC-15 (actin; see **Figure 1C-D**) and of 5H11 (PXN; see **Figure 3C-D**). Consequently, L2L can be used to correct images for inhomogeneous binding of a stain or antibody, but the restorations by a trained CNN cannot be used to quantify the distribution of the target protein in a cell. Instead, L2L could serve as image pre-processing step to extract the binary information about the location of a specific structure in a cell image, for example, of the cell membrane or the nucleus. Here, the systematic recovery of specific structure by the network might make L2L superior to classical image processing methods.

Training data for L2L can be generated with one imaging setup using two detectors simultaneously, which makes the images independent of stage drift and sample dynamics, and is time-efficient. Further, the use of unpaired training data was explored by training a cycleGAN with unaligned images of two anti-PXN antibodies (see **Figure 3A+C**). While the restored images of a cycleGAN after L2L training showed a decrease in background signal, which in the PXN dataset originated from cytosolic protein, the results were not comparable to the restorations obtained after training a CNN. We found that, since the generator network in the cycleGAN is trained to fool a discriminator based on a noisy benchmark (Y113), artefacts were introduced by the cycleGAN. This might be avoidable when training with cleaner reference images in the future.

The ability of a CNN to selectively restore specific cell structures is also highlighted in this work by training a CNN to separate a nuclear marker and plasma membrane protein label in superposed IF images (see **Figure 4**). The use of CNNs as content filter in IF microscopy could increase possibilities for multiplex imaging in the future. For example, CNNs could be employed to separate two or more markers in cell images that were acquired with microscopy setups that have a limited number of excitations sources or detectors. L2L could also be applied in multiplex imaging experiments in cases where antibodies are not selected based on performance but compatibility issues between the species in which they are raised.

Moreover, L2L could be a useful post-processing step in live cell imaging, where high-performance labels are rare. For that, the image pairs for the network training can be generated post-image acquisition *in vitro*, by fixing the cells and labelling with a higher performing antibody against the particular target structure. Our results for the caveolae dataset suggest that training a CNN with L2L data might be particularly advantageous to restore very noisy images, since it allows the implementation of a MS-SSIM loss function without introducing artefacts that we otherwise observed after training with image pairs of the same label (see **Figure 2C** and **SI Figure 3B**). Here, restored images after L2L training exhibited a high structure-to-background signal, clearly outperforming the corresponding 20 frame average images that were acquired with the STED microscope.

## Conclusion

We present a new deep learning-based image restoration method for images of cellular structures that utilises the varying performance of labels in immunofluorescence microscopy. We show that by training a CNN, that was developed for content-aware image restoration, with images of two non-identical labels that target the same cellular structure but exhibit systematic sample differences the network learns to selectively restore the correlating signal in the images. Like other methods, L2L relies on the convention of the network to under-estimate inherently unpredictable signal. However, with L2L, not only image noise but also label-induced fluorescence signal in the cell specimen can be corrected in the images after selecting appropriate training data. The ability to correct images for unspecific binding, inhomogeneous labelling of a structure or binding to cytosolic protein makes L2L, to our knowledge, unique in comparison to other deep learning-based image restoration methods that are currently used in cell biology.

## Methods

### Cell culture

For imaging, the human mesothelial cell line MeT5A (ATCC CRL-9444), the adenocarcinoma cervical cancer cell line HeLa, and the human osteosarcoma cell line U2OS were used. HeLa cells were a gift from Margaret Cunningham and U2OS cells were a gift from Kathryn McIntosh (both Strathclyde Institute of Pharmacy and Biomedical Sciences, Glasgow, UK). MeT5A cells were grown in RPMI-1640 medium (Corning), supplemented with 10% (v/v) fetal bovine serum (FBS) (Labtech), 100 μg/ml penicillin-streptomycin (Gibco), 1 mM sodium pyruvate (Gibco), 2 mM L-glutamine (Gibco) and 2 mM HEPES buffer solution (Gibco). HeLa and U2OS cells were grown in DMEM+GlutaMAX medium (Gibco) supplemented with 10% (v/v) FBS and 100 μg/ml penicillin-streptomycin. Human embryonic kidney cells HEK293T were grown in DMEM supplemented with 100 μg/ml penicillin-streptomycin and 2mM L-glutamine. All cells were kept at 37°C/5% CO_2_ in a humidified atmosphere.

### Generation and verification of knock down *PAXILLIN* MeT5A cells

Knockdown of MeT5A cells was obtained by a shRNA mediated knockdown using pKLO.1-vectors coding for shRNAs mediated targeting *PAXILLIN* (TRC N0000123137) or control. HEK293T cells were used for virus generation, and the virus was harvested, filtered and added to polybrene treated MeT5A cells. The retroviral transduction of MeT5A cells was followed by puromycin selection. Stable knockdown cell lines were verified by quantitative PCR (qPCR) (mRNA abundance) and Western blots (protein abundance). For western blotting, cell lysates were prepared in reducing lysis buffer, boiled for ten minutes and separated on conventional homemade SDS gels and developed onto Medical Blue Sensitive X-Ray Film (Scientific Laboratory Supplies). The PAXILLIN specific antibody Y113 was used (rabbit; ab32084, Abcam). Ponceau staining of the PVDF membrane was carried out to establish equal loading. For quantitative PCR (RT-qPCR), mRNA was extracted from cells using the RNeasy plus mini kit (QIAGEN). Complementary DNA (cDNA) was generated using the High-Capacity cDNA Reverse Transcriptase kit (Applied Biosystems). qPCR was performed on 1 ng of cDNA using the Brilliant III Ultra-Fast SYBR Green QPCR Master Mix (Agilent Technologies) and the Applied Biosystems QuantStudio 5 Real-Time PCR Systems. Expression of *PAXILLIN* was analysed and normalized to Hypoxanthine-guanine phosphoribosyltransferase (*HPRT1*) levels. qPCR data originate from five independent biological replicates, plotted in Prism (GraphPad) and analysed using Mann-Whitney.

### Sample preparation for immunofluorescence microscopy

If not stated otherwise, cells were plated onto #1.5 coverslips a day prior to fixation, then fixed with 4% paraformaldehyde for 15 min at 37°C, followed by a permeabilisation step with 2.5% FBS and 0.3% TX 100 in PBS for 30 min at room temperature [38]. HeLa cells that were labelled for actin were fixed 6 h after plating. The following antibodies or stains were used to generate cell specimen dual-labelled for the same target structure: to visualise the actin cytoskeleton in MeT5As the monoclonal anti-β-actin antibody AC-15 conjugated to Alexa Fluor 488 (1:250; mouse; ab6277, Abcam) and the phalloidin-Atto 565 stain (1:100; 94072, Sigma Aldrich); for α-tubulin labelling in MeT5As the two monoclonal antibodies YOL1/34 conjugated to Alexa Fluor 488 (1:500; rat; ab195883, Abcam) and DM1A (1:250; mouse; T6199, Sigma Aldrich); to label caveolae in MeT5As the monoclonal antibodies 4H312 (CAVEOLIN-1) (1:200; mouse; sc-70516, SCBT) and D1P6W (CAVIN-1) (1:200; rabbit; 69036, Cell Signal. Techn.); and for PAXILLIN labelling in MeT5A, U2OS and HeLa cells the monoclonal antibodies Y113 (1:250; rabbit; ab32084, Abcam) and 5H11 (1:500; mouse; MA5-13356, Life Technologies). Further, fixed MeT5A cells were labelled with a nuclear SYTOX Green marker (S7020, Life Technologies) and a monoclonal anti-CD44 antibody (1:100; rat; MA4400, Life Technologies). The following secondary antibodies were used: anti-rabbit Alexa Fluor 488 (1:200; donkey; A-21206, Life Technologies) to conjugate Y113; anti-rat Alexa Fluor 555 (1:100; goat; A-21434, Life Technologies) to conjugate the antibody against CD44; anti-mouse Alexa Fluor 594 (1:200; goat; A32742, Life Technologies) to conjugate 4H312; anti-mouse Alexa Fluor 633 (1:200; goat; A-11001, Life Technologies) to conjugate DM1A and 5H11; and anti-rabbit Atto 647N (1:200; goat; 40839, Sigma-Aldrich) to conjugate D1P6W. All IF samples were mounted in ProLong Glass Antifade Mountant (Life Technologies) onto microscope slides.

### Imaging

#### Confocal Microscopy

Image pairs of the IF sample labelled for the actin cytoskeleton, tubulin, PXN, as well as SYTOX Green and CD44 were taken with a commercial confocal laser scanning microscope (Leica TCS SP8, Leica Microsystems), using a HC PL Apo 63x/1.4 N.A. CS2 objective with no digital zoom (for image sizes see **Table 2**). To acquire data for L2L training, both markers in the individual sample were imaged simultaneously using the two in-built photomultiplier detectors, each equipped with a prism-based tunable spectral filter. Each field of view was imaged twice to acquire the two noise realisations of the sample that were used for N2N training of a CNN. The sample dual-labelled for actin was excited with a 488 nm and 552 nm laser line and the two spectral detectors were set to detect light between 495-540 and 560-700 nm. The samples dual-labelled for tubulin and PXN were excited with a 488 nm and 638 nm laser line and the two spectral detectors were set to detect light between 495-620 nm and 645-750 nm. The IF sample labelled for SYTOX Green and CD44 was imaged by setting the spectral range of the detector to 560-660 nm. Separate images of SYTOX and CD44 were acquired exciting the sample with a 488 and 552 nm laser line, respectively. A corresponding, superposed image of both markers was acquired by exciting the sample with both lasers simultaneously.

**Table 2.** Training settings. Overview of the training data and settings in the CSBDeep framework for L2L training.

| Dataset | Actin | Tubulin | Caveolae | PAXILLIN | SYTOX+CD44 |
| --- | --- | --- | --- | --- | --- |
| Input marker | AC-15 | DM1A | D1P6W | 5H11 | *superposed*<sup>‡</sup> |
| Benchmark marker | Phalloidin | YOL1/34 | 4H312 | Y113 | SYTOX/CD44<sup>‡</sup> |
| No. of raw image pairs | 68+272<sup>†</sup> | 51+204<sup>†</sup> | 60+240<sup>†</sup> | 77+308<sup>†</sup> | 58 |
| Size of raw images | (4096 px)<sup>2</sup> | (4096 px)<sup>2</sup> | (7730 px)<sup>2</sup> | (4096 px)<sup>2</sup> | (2608 px)<sup>2</sup> |
| Percentiles for normalisation | 1/99.9 | 1/99.9 | 2/99.5 | 1/99.9 | 2/99.8 |
| No. of validation /<br>training patches (128 px)<sup>2</sup> | 13,926 /<br>125,338 | 10,445 /<br>94,003 | 18,432 /<br>165,888 | 15,770 /<br>141,926 | 11,878 /<br>106,906 |
| Learning rate (LR) | 2E-4 | 2E-4 | 1E-5 | 2E-4 | 2E-5 |
| Batch size / steps per epoch /<br>epoch no. | 32 / 100 /<br>400 | 32 / 100 /<br>500 | 16 / 500 /<br>100 | 32 / 100 /<br>500 | 32 / 1000 /<br>250 |

<sup>†</sup> augmented images, <sup>‡</sup> two channel images

#### Stimulated emission depletion microscopy

Fixed MeT5A cells dual-labelled for tubulin were further imaged with a Leica SP8 TCS 3X STED microscope, using a Leica HC PL APO 100x/1.40 Oil STED WHITE objective (Leica 15506378). The sample was excited with a Supercontinuum White Light Laser at 488 nm and 633 nm for confocal and at 633 nm for STED imaging. For confocal imaging, two Leica PMT detectors were set to detect light between 498-600 nm and 643-743 nm. For STED imaging, a Leica HyD hybrid detector was set to detect light between 643-743nm. STED depletion was performed with a Leica 775 nm depletion laser set to 50% with time gating from 0.3-8 ns. Pairs of confocal and STED images were acquired with a 15 nm pixel size. Sequential STED image pairs of the cell specimen labelled for caveolae were acquired by first exciting the sample at 646 nm and detecting light between 656-750 nm, then exciting at 591 nm and using a spectral range of 601-650 nm for the detection. The depletion laser power was set to 100% with time gating from 0.3-8 ns, and the pixel size was set to 10 nm.

### Image drift correction

Fluorescence images that were taken sequentially were corrected for stage drift by estimating the extent of the displacement between the images. 100 image pairs of size 60 pixels x 60 pixels were generated for each image in a temporal image stack, after applying a Gaussian filter (*sigma*=2) and using the first detected image as benchmark. For that, the *create_patches* function from the CSBDeep framework was used that cuts random image patch pairs in non-background areas of the image. For each created image patch, the SSIM index (filter size=11, *sigma*=1.5) was calculated by shuffling the uncorrected image patch relative to the benchmark patch by 5 pixels into all directions, effectively cutting both into 50 px x 50 px patches. The shuffling vector that, on average, yielded the highest structural similarity between both images was selected as optimal position for the sequential image, and the images were cropped accordingly.

### Data augmentation and pre-processing

Images were augmented to create more training data. For that, the images were interpolated, using the zooming factors 0.5, 0.75, 1.25 and 1.5 (0.8, 0.9, 1.1 and 1.2 for the caveolae dataset), and then randomly rotated by 0°, 90°, 180° or 270°. STED images for the caveolae dataset were pre-processed by adding Poisson noise to broaden the image histogram that, due to being acquired with hybrid detectors, was nearly binary in the raw images, resulting in poor results for the network trainings. Prior to the training patch generation in the CSBDeep framework, a percentile normalisation was conducted (see **Table 2**).

### Training the convolutional neural network

For N2N and L2L training, the CSBDeep framework was used, which is a CNN with U-Net architecture [2] that was developed for content-aware image restoration in fluorescence microscopy [10]. The CSBDeep framework (version 0.6.0) was downloaded from github and used with its default settings unless stated otherwise (https://github.com/CSBDeep/CSBDeep).

In the CSBDeep framework, training patches were generated randomly from the raw images after pre-processing (see previous section). Half of the training patches were generated from the raw images, the other half from augmented images. Validation data was generated via a 90/10 train/test split, and the training and validation loss were monitored to rule out overfitting.

A least absolute deviation loss function (*L*_*1*_) and different multi-scale SSIM loss functions were used for the trainings (*L*_*SSIM*_ (*M*=1), *L*_*3S-SSIM*_ (*M*=3), *L*_*5S-SSIM*_ (*M*=5)). For a *L*_*3S-SSIM*_, the weights were set to (0.2096, 0.4659, 0.3245), for a *L*_*5S-SSIM*_, the filter size for the Gaussian filter was set to 7; otherwise the suggested settings in [35] were used.

The settings that were applied for the L2L trainings are shown in **Table 2**. The same settings were used to train the network for N2N, but the training data was generated from two sequential images of the respective antibody.

To train the CNN as separator of two cellular markers (the SYTOX Green stain and a CD44 antibody) in superposed IF images, the CSBDeep Framework was trained with two channel images, where the input consisted of the superposed image in both channels, and the separately acquired images of SYTOX and CD44 as the output channels (see **Table 2** (r*ight*)).

To further evaluate L2L, repeated 8- or 10-fold cross validations were conducted on the datasets using the settings as outlined in **Table 3**. The selection of the raw image pairs from the total dataset for each cross validation and the fold allocation were conducted randomly in Python. The image pairs for each training were generated as described above, including the pre-processing, and disabling the train/test split in the CSBDeep Framework. The relative change of the *NRMSE* and *5S-SSIM* index were calculated between input/benchmark and prediction/benchmark for the test images of each fold, deriving an average for each cross-validation (see **Figure 5**).

**Table 3.**
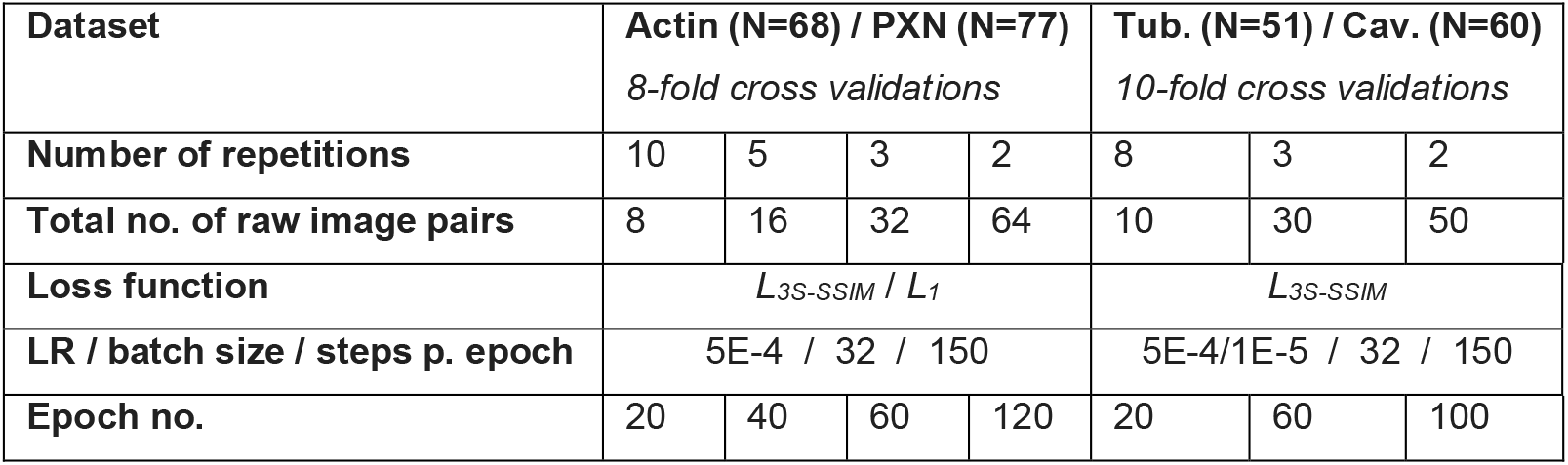
Settings for repeated cross validations. Overview of the settings in the CSBDeep framework that were used for repeated 8- or 10-fold cross validations, training the network for L2L with the different datasets.

### Training the cycle generative adversarial network

The implementation of a CycleGAN in Pytorch was downloaded from github and, if not stated otherwise, trained with the default parameters (https://github.com/junyanz/pytorch-CycleGAN-and-pix2pix) [13]. The CycleGAN was trained with unaligned images of the PXN dataset that were pre-processed as outlined in the previous sections (see also **Table 2**), using a Least Squares GAN (LSGAN) with a ResNet-9 generator architecture and a 70×70 PatchGAN discriminator architecture. Training was conducted with a batch size of 4, an epoch number of 4 (3 with linear decay of the learning rate) and a scaling factor of 0.0005 for the network initialization.

## Contributions

L.S.K. designed the method, prepared the samples for imaging, generated the imaging data and wrote/implemented Python scripts for pre-processing, trainings and evaluations. C.G.H. and O.S. developed the *PXN* knockdown cells. O.S. validated the knockdown. J.V. and L.S.K. took the confocal-STED image pairs. L.S.K., C.G.H. and G.M. wrote the manuscript, with input from all authors.

## Acknowledgements

L.S.K. was supported by the Medical Research Council and Engineering and Physical Sciences Research Council Centre for Doctoral Training in Optical Medical Imaging (Optima), grant number EP/L016559/1. J.V. was supported by the Wellcome Trust, grant number 208345/Z/17/Z. STED imaging was performed at the Edinburgh Super Resolution Imaging Consortium (ESRIC), which is supported by the MRC and the Wellcome Trust. Work on-going in the Gram Hansen lab is supported by a University of Edinburgh Chancellor’s Fellowship, as well as by the Worldwide Cancer Research and the June Hancock Mesothelioma Research Fund (JHMRF). G.M. was supported by the grants MR/K015583/1, BB/P02565X/1 and BB/T011602/1. The authors thank Martin Weigert (Swiss Federal Institute of Technology Lausanne, Switzerland) for his helpful feedback.

## Supplementary Information

**SI Figure 1.**
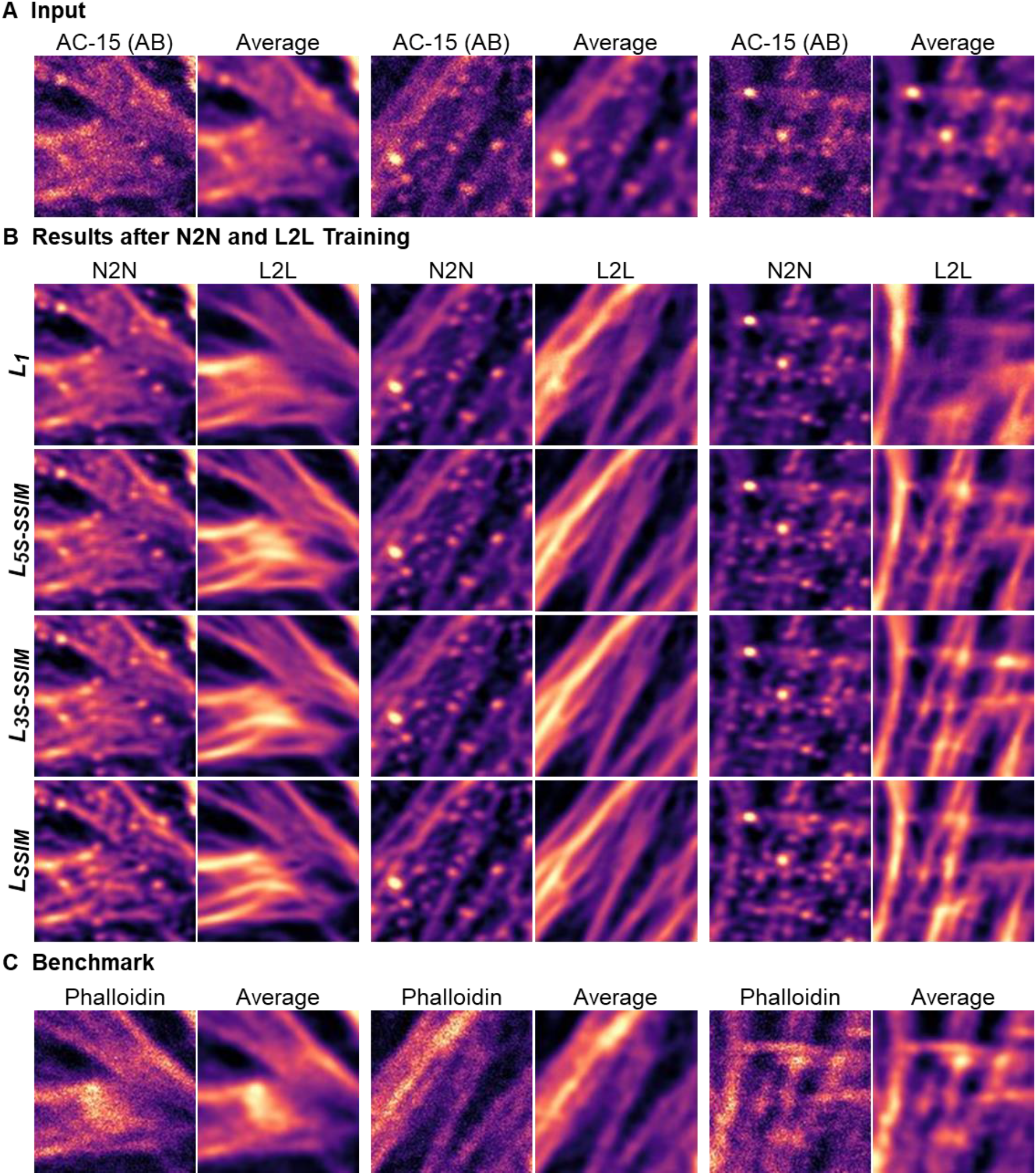
Loss function-dependent label2label (L2L) and noise2noise (N2N) results for images of actin. Three representative examples of confocal image pairs of HeLa cells that were dual-labelled with the anti-β-actin antibody (AB) AC-15 and a phalloidin-Atto 565 stain. (**A**) Images of AC-15 in a HeLa cell and corresponding 20-frame average images. (**B**) Restored images of AC-15 by a CNN after N2N and L2L training, dependent on the training loss function. For N2N training, two sequential images of AC-15 in HeLa cells were used as training input and benchmark. For L2L training, benchmark images were replaced by (**C**) cell images of a phalloidin stain. The shown images were excluded from the trainings and contrast adjusted to the 2^nd^ and 99.8^th^ percentile (image dimensions: 4.5 µm x 4.5 µm).

**SI Figure 2.**
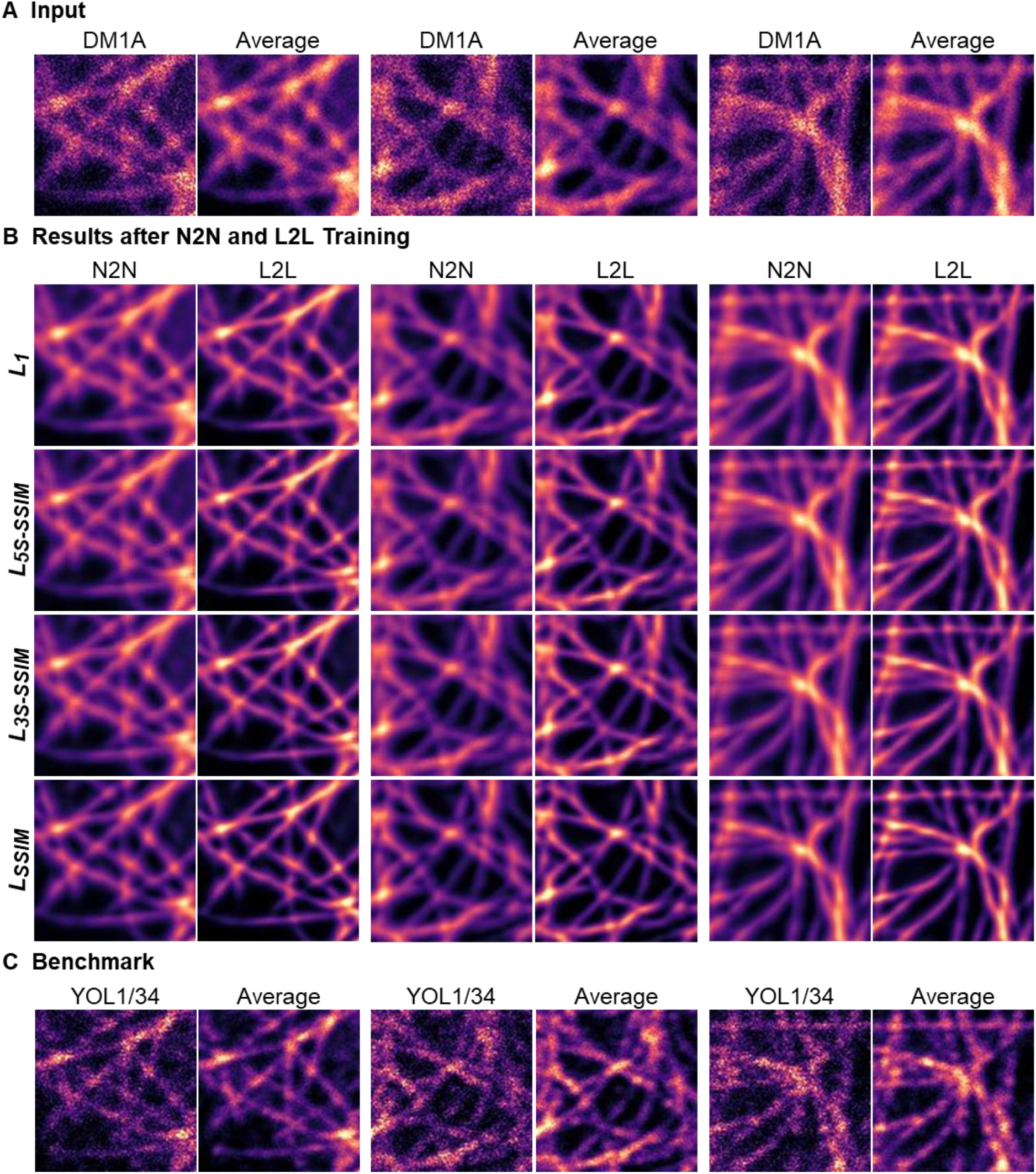
Loss function-dependent label2label (L2L) and noise2noise (N2N) results for images of tubulin. Three representative examples of confocal image pairs of MeT5A cells that were dual-labelled with the anti-α-tubulin antibodies DM1A and YOL1/34. (**A**) Images of DM1A in MeT5A cells and corresponding 20-frame average images. (**B**) Restored images of DM1A in MeT5A cells by a CNN after N2N and L2L training, dependent on the training loss function. For N2N training, two sequential images of the DM1A signal in MeT5A cells were used as training input and benchmark. For L2L training, benchmark images were replaced by (**C**) cell images of the anti-α-tubulin antibody YOL1/34. The shown images were excluded from the trainings and contrast adjusted to the 2^nd^ and 99.8^th^ percentile (image dimensions: 4.5 µm x 4.5 µm).

**SI Figure 3.**
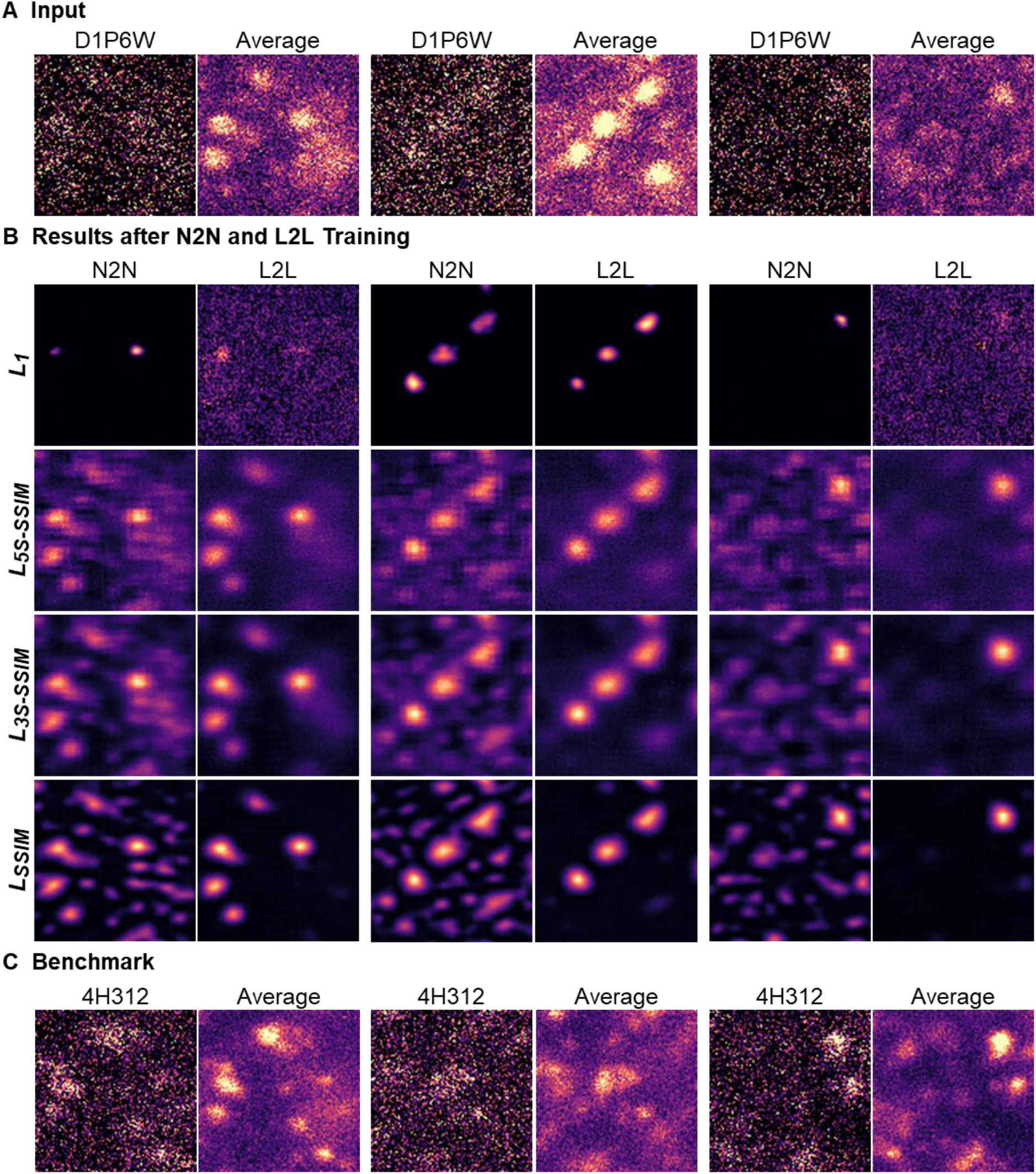
Loss function-dependent label2label (L2L) and noise2noise (N2N) results for images of caveolae. Three representative examples of STED image pairs of MeT5A cells that were dual-labelled with the anti-CAVIN-1 antibody D1P6W and anti-CAVEOLIN-1 antibody 4H312. (**A**) Images of D1P6W in MeT5A cells and corresponding 20-frame average images. (**B**) Restored images of D1P6W in MeT5A cells by a CNN after N2N and L2L training, dependent on the training loss function. For N2N training, two sequential images of the D1P6W signal in MeT5A cells were used as training input and benchmark. For L2L training, benchmark images were replaced by (**C**) cell images of the antibody 4H312. The shown images were excluded from the trainings and contrast adjusted to the 2^nd^ and 100^th^ percentile (image dimensions: 1 µm x 1 µm).

**SI Figure 4.**
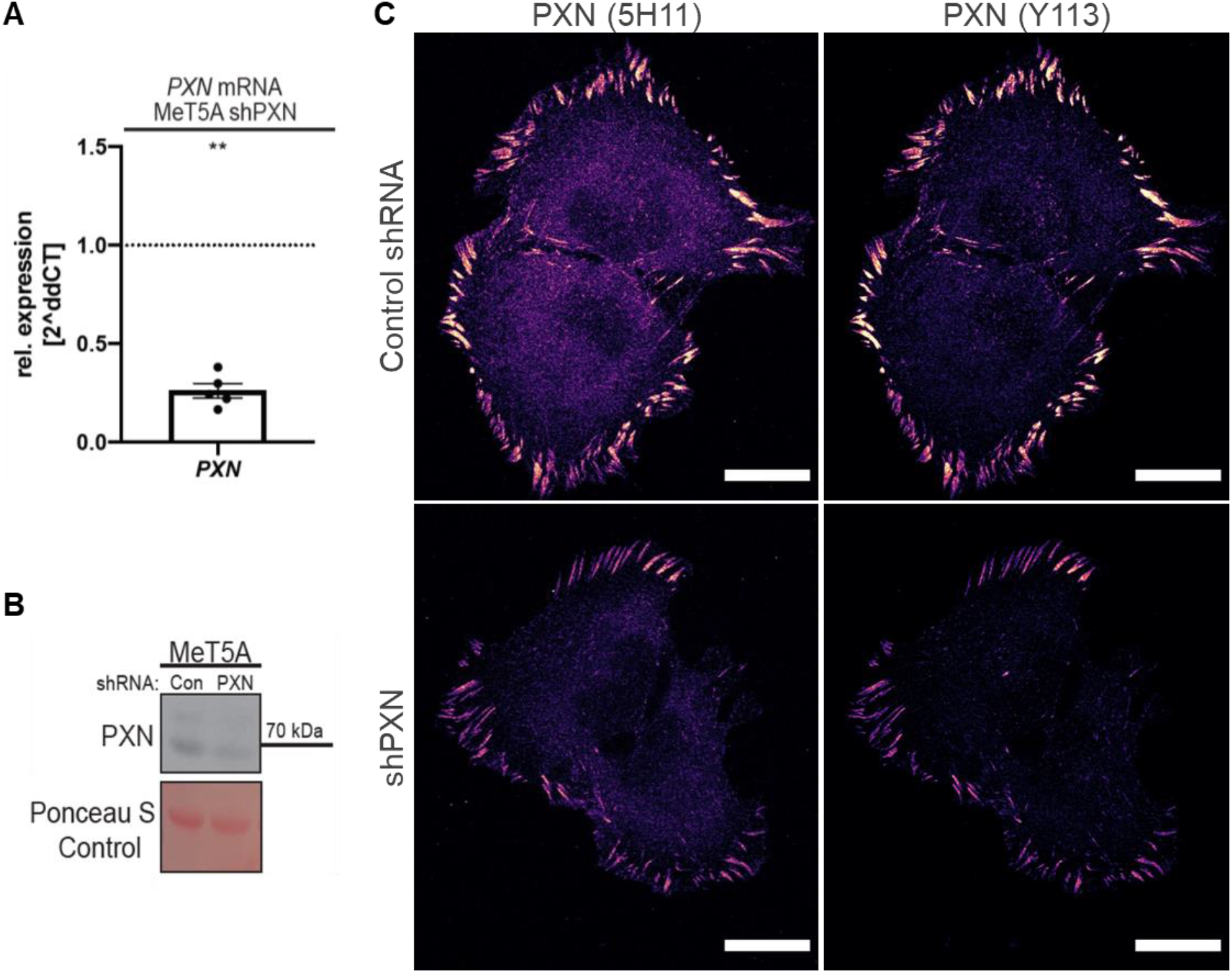
*PAXILLIN* (*PXN*) knockdown in MeT5A cells. PXN is a focal adhesion molecule. (**A**) Results of a RT-qPCR analysis, including the standard error of the mean: the relative expression of the *PXN* knockdown cells is 26.14±0.04%. (**B**) Western analysis of cell lysates of MeT5A cells visualising the PXN protein knockdown, with Ponceau stain ensuring equal loading. The monoclonal anti-PXN antibody Y113 was used. The PXN expression is reduced by around a half in the knockdown cells compared to the control. (**C**) Immunofluorescence images of MeT5A cells that were dual-labelled with the anti-PXN antibodies (*left*) 5H11 and (*right*) Y113 after (*top*) mock-infection and (*bottom*) shRNA mediated *PXN*-knockdown. The low signal of cytosolic protein in the knockdown cells for both antibodies indicate that the difference in cytosolic signal between the two, as observed in the control cells, originates from clone-dependent antibody binding to cytosolic PXN rather than unspecific binding (scale bar = 20 µm).

**SI Figure 5.**
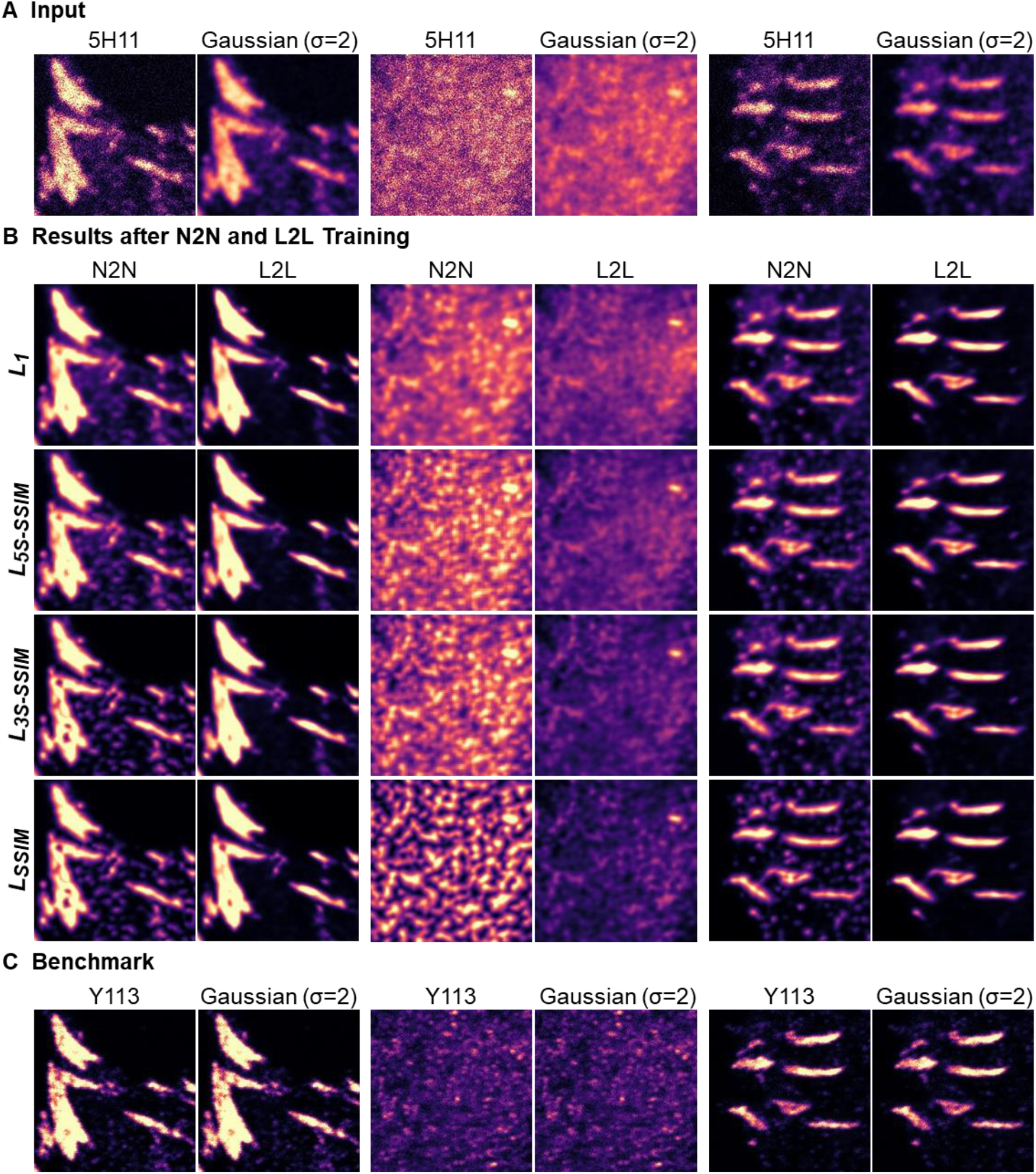
Loss function-dependent label2label (L2L) and noise2noise (N2N) results for images of focal adhesions. Three representative examples of confocal image pairs of MeT5A cells that were dual-labelled with the anti-PAXILLIN (PXN) antibodies 5H11 and Y113. (**A**) Images of 5H11 in MeT5A cells and corresponding Gaussian filter images (s=2). (**B**) Restored images of 5H11 in a MeT5A cell by a CNN after N2N and L2L training, dependent on the training loss function. For N2N training, two sequential images of 5H11 in MeT5A cells were used as training input and benchmark. For L2L training, benchmark images were replaced by (**C**) cell images of the anti-PXN antibody Y113. The shown images were excluded from the trainings. Images of 5H11 and Y113 were contrast adjusted to the 2^nd^ and 99.8^th^ percentile of the respective original image (image dimensions: 9.1 µm x 9.1 µm).

## References

[1] C. Belthangady and L. A. Royer, “Applications, promises, and pitfalls of deep learning for fluorescence image reconstruction,” Nat. Methods, vol. 16, no. 12, pp. 1215–1225, Dec. 2019, doi: 10.1038/s41592-019-0458-z.

[2] O. Ronneberger, P. Fischer, and T. Brox, “U-Net: Convolutional Networks for Biomedical Image Segmentation,” Med. Image Comput. Comput. Interv. – MICCAI 2015. MICCAI 2015. Lect. Notes Comput. Sci., vol. 9351, pp. 234–241, 2015, doi: 10.1007/978-3-319-24574-4_28.

[3] C. Ounkomol, S. Seshamani, M. M. Maleckar, F. Collman, and G. R. Johnson, “Label-free prediction of three-dimensional fluorescence images from transmitted-light microscopy,” Nat. Methods, vol. 15, no. 11, pp. 917–920, Nov. 2018, doi: 10.1038/s41592-018-0111-2.

[4] E. M. Christiansen et al., “In Silico Labeling: Predicting Fluorescent Labels in Unlabeled Images,” Cell, vol. 173, no. 3, pp. 792–803, Apr. 2018, doi: 10.1016/J.CELL.2018.03.040.

[5] T. Falk et al., “U-Net: deep learning for cell counting, detection, and morphometry,” Nat. Methods, vol. 16, no. 1, pp. 67–70, Jan. 2019, doi: 10.1038/s41592-018-0261-2.

[6] J. C. Caicedo et al., “Evaluation of Deep Learning Strategies for Nucleus Segmentation in Fluorescence Images,” Cytom. Part A, vol. 95, no. 9, pp. 952–965, Sep. 2019, doi: 10.1002/cyto.a.23863.

[7] M. C. Daniel et al., “Automated segmentation of the corneal endothelium in a large set of ‘real-world’ specular microscopy images using the U-Net architecture,” Sci. Rep., vol. 9, p. 4752, Dec. 2019, doi: 10.1038/s41598-019-41034-2.

[8] I. Jansen et al., “Automated Detection and Grading of Non-Muscle-Invasive Urothelial Cell Carcinoma of the Bladder.,” Am. J. Pathol., vol. 190, no. 7, pp. 1483–1490, Jul. 2020, doi: 10.1016/j.ajpath.2020.03.013.

[9] T. Suzuki, T. Matsuzaki, H. Hagiwara, T. Aoki, and K. Takata, “Recent advances in fluorescent labeling techniques for fluorescence microscopy,” Acta Histochemica et Cytochemica, vol. 40, no. 5. Japan Society of Histochemistry and Cytochemistry, pp. 131–139, 2007, doi: 10.1267/ahc.07023.

[10] M. Weigert et al., “Content-aware image restoration: pushing the limits of fluorescence microscopy,” Nat. Methods, vol. 15, no. 12, pp. 1090–1097, Dec. 2018, doi: 10.1038/s41592-018-0216-7.

[11] I. J. Goodfellow et al., “Generative adversarial nets,” in Advances in Neural Information Processing Systems, 2014, vol. 3, no. January, pp. 2672–2680, doi: 10.3156/jsoft.29.5_177_2.

[12] H. Wang et al., “Deep learning enables cross-modality super-resolution in fluorescence microscopy,” Nat. Methods, vol. 16, no. 1, pp. 103–110, Jan. 2019, doi: 10.1038/s41592-018-0239-0.

[13] J. Y. Zhu, T. Park, P. Isola, and A. A. Efros, “Unpaired Image-to-Image Translation Using Cycle-Consistent Adversarial Networks,” in Proceedings of the IEEE International Conference on Computer Vision, Dec. 2017, pp. 2242–2251, doi: 10.1109/ICCV.2017.244.

[14] L. von Chamier et al., “Democratising deep learning for microscopy with ZeroCostDL4Mic,” Nat. Commun., vol. 12, no. 1, p. 2276, Dec. 2021, doi: 10.1038/s41467-021-22518-0.

[15] S. Lim, S. E. Lee, S. Chang, B. Sim, and J. C. Ye, “CycleGAN with a blur kernel for deconvolution microscopy: Optimal transport geometry,” IEEE Trans. Comput. IMAGING, vol. 6, pp. 1127–1138, 2020.

[16] A. Krull, T.-O. Buchholz, and F. Jug, “Noise2Void - Learning Denoising from Single Noisy Images,” Proc. IEEE Conf. Comput. Vis. Pattern Recognit., pp. 2129–2137, 2019, doi: 10.1109/CVPR.2019.00223.

[17] M. Prakash, A. Krull, and F. Jug, “DivNoising: Diversity Denoising with Fully Convolutional Variational Autoencoders,” arXiv Prepr., 2020, Accessed: Jul. 24, 2020. [Online]. Available: http://arxiv.org/abs/2006.06072.

[18] J. Lehtinen et al., “Noise2Noise: Learning Image Restoration without Clean Data,” Proc. 35th Int. Conf. Mach. Learn., vol. 80, pp. 2965–2974, Jul. 2018.

[19] C. B. Saper, “A guide to the perplexed on the specificity of antibodies,” J. Histochem. Cytochem., vol. 57, no. 1, pp. 1–5, 2009, doi: 10.1369/jhc.2008.952770.

[20] D. M. Miller and D. C. Shakes, “Immunofluorescence Microscopy,” Methods Cell Biol., vol. 48, pp. 365–394, Jan. 1995, doi: 10.1016/S0091-679X(08)61396-5.

[21] C. Wagner et al., “Dynamic force spectroscopy on the binding of monoclonal antibodies and tau peptides,” Soft Matter, vol. 7, no. 9, pp. 4370–4378, May 2011, doi: 10.1039/c0sm01414a.

[22] R. Dominguez and K. C. Holmes, “Actin Structure and Function,” Annu. Rev. Biophys., vol. 40, no. 1, pp. 169–186, Jun. 2011, doi: 10.1146/annurev-biophys-042910-155359.

[23] P. Vedula et al., “Diverse functions of homologous actin isoforms are defined by their nucleotide, rather than their amino acid sequence,” Elife, vol. 6, Dec. 2017, doi: 10.7554/eLife.31661.

[24] C. Suarez and D. R. Kovar, “Internetwork competition for monomers governs actin cytoskeleton organization,” Nature Reviews Molecular Cell Biology, vol. 17, no. 12. 2016, doi: 10.1038/nrm.2016.106.

[25] V. Dugina, I. Zwaenepoel, G. Gabbiani, S. Clément, and C. Chaponnier, “Beta and gamma-cytoplasmic actins display distinct distribution and functional diversity.,” J. Cell Sci., vol. 122, pp. 2980–2988, Aug. 2009, doi: 10.1242/jcs.041970.

[26] E. Nogales, “Structural insights into microtubule function,” Annual Review of Biochemistry, vol. 69. pp. 277–302, Nov. 2000, doi: 10.1146/annurev.biochem.69.1.277.

[27] F. Pellegrini and D. R. Budman, “Review: Tubulin function, action of antitubulin drugs, and new drug development,” Cancer Investigation, vol. 23, no. 3. pp. 264–273, 2005, doi: 10.1081/CNV-200055970.

[28] C. G. Hansen and B. J. Nichols, “Exploring the caves: Cavins, caveolins and caveolae,” Trends in Cell Biology, vol. 20, no. 4. Elsevier, pp. 177–186, Apr. 01, 2010, doi: 10.1016/j.tcb.2010.01.005.

[29] I. M. Khater, F. Meng, T. H. Wong, I. R. Nabi, and G. Hamarneh, “Super Resolution Network Analysis Defines the Molecular Architecture of Caveolae and Caveolin-1 Scaffolds,” Sci. Rep., vol. 8, no. 1, pp. 1–15, Dec. 2018, doi: 10.1038/s41598-018-27216-4.

[30] V. Rausch and C. G. Hansen, “The Hippo Pathway, YAP/TAZ, and the Plasma Membrane,” Trends in Cell Biology, vol. 30, no. 1. Elsevier Ltd, pp. 32–48, Jan. 01, 2020, doi: 10.1016/j.tcb.2019.10.005.

[31] F. Martino, A. R. Perestrelo, V. Vinarský, S. Pagliari, and G. Forte, “Cellular mechanotransduction: From tension to function,” Frontiers in Physiology, vol. 9. Frontiers Media S.A., p. 824, Jul. 05, 2018, doi: 10.3389/fphys.2018.00824.

[32] H. Zhao, O. Gallo, I. Frosio, and J. Kautz, “Loss Functions for Image Restoration with Neural Networks,” IEEE Trans. Comput. IMAGING, vol. 3, no. 1, 2017.

[33] A. B. L. Larsen, S. K. Sønderby, H. Larochelle, and O. Winther, “Autoencoding beyond pixels using a learned similarity metric,” Proc. 33rd Int. Conf. Mach. Learn., no. 48, pp. 1558–1566, 2016.

[34] Z. Wang, A. C. Bovik, H. Rahim Sheikh, and E. P. Simoncelli, “Image Quality Assessment: From Error Visibility to Structural Similarity,” IEEE Trans. IMAGE Process., vol. 13, no. 4, 2004, doi: 10.1109/TIP.2003.819861.

[35] Z. Wang, E. P. Simoncelli, and A. C. Bovik, “Multi-scale structural similarity for image quality assessment,” Conf. Rec. Asilomar Conf. Signals, Syst. Comput., vol. 2, pp. 1398–1402, 2003, doi: 10.1109/acssc.2003.1292216.

[36] E. Peli, “Contrast in complex images,” J. Opt. Soc. Am. A, vol. 7, no. 10, pp. 2032–2040, 1990, doi: 10.1364/josaa.7.002032.

[37] M. Bates, B. Huang, G. T. Dempsey, and X. Zhuang, “Multicolor super-resolution imaging with photo-switchable fluorescent probes,” Science, vol. 317, no. 5845, pp. 1749–1753, Sep. 2007, doi: 10.1126/science.1146598.

[38] V. Rausch and C. G. Hansen, “Immunofluorescence Study of Endogenous YAP in Mammalian Cells,” Humana Press, New York, NY, 2019, pp. 97–106.

